# Chromosome evolution in Lepidoptera

**DOI:** 10.1101/2023.05.12.540473

**Authors:** Charlotte J. Wright, Lewis Stevens, Alexander Mackintosh, Mara Lawniczak, Mark Blaxter

## Abstract

Chromosomes are a central unit of genome organisation. One tenth of all described species on Earth are Lepidoptera, butterflies and moths, and these generally possess 31 holocentric chromosomes. However, a subset of lepidopteran species display dramatic variation in chromosome counts. By analysing 210 chromosomally-complete lepidopteran genomes, the largest analysis of eukaryotic chromosomal-level reference genomes to date, we show that the diverse karyotypes of extant species are derived from 32 ancestral linkage groups, which we term Merian elements. Merian elements have remained largely intact across 250 million years of evolution and diversification. Against this stable background, we identify eight independent lineages that have evaded constraint and undergone extensive reorganisation - either by numerous fissions or a combination of fusion and fission events. Outside these lineages, fusions are rare and fissions are rarer still. Fusions tend to involve small, repeat-rich Merian elements and/or the Z chromosome. Together, our results reveal the constraints on genome architecture in Lepidoptera and enable a deeper understanding of the importance of chromosomal rearrangements in shaping the evolution of eukaryotic genomes.

---

Chromosomes, the central units of genome architecture in eukaryotic organisms, are key to processes including recombination and segregation. While chromosomes are generally stable over evolutionary time, large-scale rearrangements, such as fusions and fissions can occur. Consequently, the chromosomes of extant species can be used to infer the linkage groups present in a common ancestor, termed ancestral linkage groups (ALGs). ALGs have been identified in many taxa including Diptera (Bhutkar *et al*., 2008), flowering plants (Murat *et al*., 2017), Nematoda (Tandonnet *et al*., 2019; Gonzalez de la Rosa *et al*., 2021), mammals (Band *et al*., 2000), vertebrates (Simakov *et al*., 2020) and Metazoa (Simakov *et al*., 2022). Chromosomal rearrangements have important consequences for genome function (Marquès-Bonet *et al*., 2004), speciation (Feulner and De-Kayne, 2017), and adaptation (De Storme and Mason, 2014). For example, heterozygous chromosomal fusions can interfere with meiosis, resulting in reproductively isolated populations (White, 1968; Hauffe and Searle, 1998). The evolutionary forces that constrain chromosome number and maintain ALGs remain unclear. Moreover, how and why certain taxa evade such constraints, resulting in a high rate of karyotypic change, is not understood.

In monocentric chromosomes a single region, the centromere, serves as the organising centre for Mendelian partitioning of homologues during mitosis and meiosis. Discrete centromeres are absent in the chromosomes of holocentric taxa, where centromeric functions are dispersed along the chromosome. Holocentricity has evolved independently at least nine times in animals, including nematodes, several times in arthropods, and four times in plants (Schrader, 1935; Albertson and Thomson, 1982; Benavente, 1982; Luceño, Vanzela and Guerra, 1998; Kuta *et al*., 2004; Melters *et al*., 2012). The most speciose of these holocentric groups is Amphiesmenoptera, comprising the insect orders Lepidoptera (moths and butterflies) and Trichoptera (caddisflies), which together account for 15% of all described eukaryotic species (Mallet, 2011; Morse, 2011). The convergent evolution of holocentricity in multiple speciose groups indicates that this alternative solution to accurate segregation of chromosomes may be evolutionarily advantageous.

Holocentric chromosomes are hypothesised to facilitate rapid karyotypic evolution as fragments derived from fission maintain kinetochore function (Marec *et al*., 2001; Lukhtanov *et al*., 2018). However, chromosome numbers and their gene content are generally stable over evolutionary time in both holocentric and monocentric taxa (Ruckman *et al*., 2020). In Lepidoptera, most species have a haploid chromosome number (hereafter chromosome number, *n*) of *n*=29-31 (Robinson, 1971; White, 1977). However, some groups exhibit dramatic variation in chromosome number, with haploid counts ranging from 5 to 223 (Brown, Von Schoultz and Suomalainen, 2004; Lukhtanov, 2015). Lepidoptera are thus the most karyotypically diverse group of any non-polyploid eukaryote (Lukhtanov, 2015; Hill *et al*., 2019). While holocentricity may make chromosomal variation more likely to evolve, the overwhelming conservation of chromosome number observed in Lepidoptera suggests that additional mechanisms must constrain karyotypic evolution.

Changes in chromosome number alter the recombination rate (Dumas and Britton-Davidian, 2002; Butlin, 2005). In Lepidoptera, there tends to be one crossover event per chromosome per generation (Kaback *et al*., 1992; Jiggins *et al*., 2005; Yasukochi *et al*., 2006), and thus loci on a fused chromosome formed from two equally-sized progenitors will experience a per base recombination rate of approximately half what they would have experienced in the unfused state. Changes in recombination rate will impact the evolutionary forces that shape genome architecture, altering the effect of selection at linked sites and thus the effective population size. Lower recombination rates may also intensify Hill-Robertson interference between tightly-linked beneficial loci, hindering adaptive evolution (Hill and Robertson, 1966). However, local adaptation is facilitated by reduced recombination between locally adapted loci in the presence of gene flow (Charlesworth and Charlesworth, 1979; Lenormand and Otto, 2000).

Here, we infer the ALGs of Lepidoptera, which we term Merian elements, using a reference-free, phylogenetically-aware approach with 210 chromosomal genome assemblies - the largest set of chromosomal lepidopteran genomes analysed to date. We find that Merian elements have remained largely intact in most species. While infrequent fusions occur, fissions are extremely rare. Constraints on large-scale reorganisation have been relaxed in eight lineages, resulting in either many fissions or numerous fusion and fission events. Across Lepidoptera, we find that fusions are biased towards the shorter autosomes and the sex chromosome, suggesting that both chromosome length and haploidy in the heterogametic sex play key roles in shaping the constraints on genomic change.

## Results

### Over two hundred chromosomally-complete lepidopteran genomes

To explore karyotype variation across Lepidoptera, we selected chromosome-level reference genomes for 210 species of Lepidoptera, representing 16 of the 43 (37%) superfamilies, including basal lineages such as Micropterigidae and Tineidae. Almost 90% of the assemblies (188 of 210) were generated by the Darwin Tree of Life project and therefore derive from taxa present in Britain and Ireland (table S1). The lepidopteran reference genomes have high biological completeness (mean 98.24%, sd=1.75%); assessed by Benchmarking Using Single Copy Orthologs (lepidoptera odb10 dataset), high contiguity (mean contig N50 13.47, sd=6.92 Mb) and the vast majority of each assembly is scaffolded into chromosomes (mean 99.56%, sd=1.28) (table S1, S3, fig. S1, fig. S2).

We inferred the phylogeny of the sampled species using 4,947 single-copy orthologues present in 90% of all species (Fig. 1A) (Simão *et al*., 2015), which we rooted using five representative species of the two main suborders from the sister group, Trichoptera (caddisflies). Four-fifths (82%) of the lepidopteran species had an assembled *n* of 28-31 (Fig. 1B, fig. S3). The karyotypes inferred from the genome assemblies are consistent with previous cytological determinations, with low numbers in *Brenthis* (e.g. *n*=14 in *Brenthis ino*) and high numbers in *Lysandra* and *Leptidea* (e.g. *n*=90 in *Lysandra coridon*) (Robinson, 1971; de Vos *et al*., 2020). Genome size varied tenfold, from 230 Mb in *Aporia crataegi* to 2.29 Gb in *Euclidia mi* (Fig. 1C, fig. S4). In contrast to previous studies (de Vos *et al*., 2020), we found no significant correlation between genome size and chromosome number (fig. S5).

**Fig. 1:**
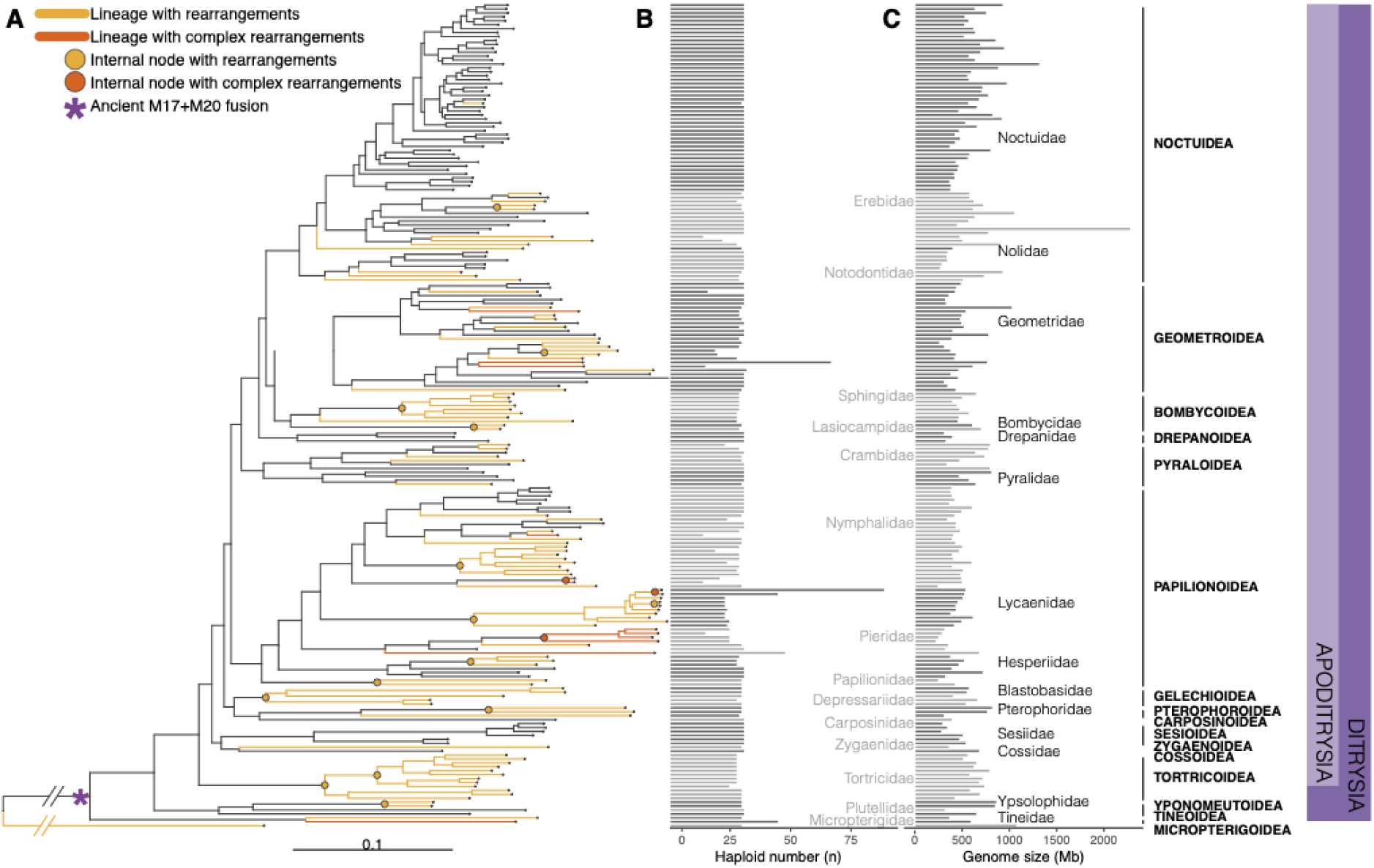
Phylogenetic relationships of 210 lepidopteran species and the distribution of large-scale rearrangement events. (A) Phylogeny inferred using IQ-TREE with a supermatrix of 4,947 orthologues under the LG substitution model with gamma-distributed rate variation among sites. Half of the species have retained 31 intact Merian elements since the last common ancestor of Lepidoptera (black lines). Orange branches indicate lineages with at least one fusion or fission event. Orange circles indicate internal nodes where descendents share a fusion event. We inferred no fission events at internal nodes. Red branches indicate lineages with extensively reorganised genomes (*Lysandra coridon, Lysandra bellargus, Pieris brassicae, Pieris napi, Pieris rapae, Tinea semiifulvella, Melinaea menophilus, Melinaea marsaeus, Aporia crataegi, Brenthis ino, Operophtera brumata, Philereme vetulata, Leptidea sinapis* and *Apeira syringaria)*. The distribution of (B) chromosome number and (C) genome size (Mb) across 210 lepidopteran species. Alternating shades distinguish different taxonomic families.

We observed strong patterning of features along each chromosome, including GC content, repeat and coding densities, consistent with observations in *Heliconius* butterflies (Cicconardi et al., 2021). Both GC content and the density of repetitive sequences were higher towards the ends of chromosomes compared to the centre (fig. S6A,B). In contrast, coding density tended to decrease at the ends of chromosomes and increase towards the centre of chromosomes (fig. S6C). Normalising for chromosome length, we found that the pattern of feature distribution was similar across all autosomes (fig. S6D). Z chromosomes had the same pattern as autosomes (fig. S6D).

### Thirty-two ancestral lepidopteran linkage groups

We used 5,287 single-copy orthologues in 210 lepidopteran and 4 trichopteran species to define ALGs in a reference-free, phylogenetically-aware manner, using the tool Syngraph, described in (Mackintosh *et al*., 2023). Although previous work proposed 31 ALGs in the last common ancestor of Lepidoptera (d’Alençon *et al*., 2010; Heliconius Genome Consortium, 2012; Ahola *et al*., 2014), we assigned 4,112 orthologues (78%) to 32 ALGs (Fig. 2A): 31 autosomes and Z, the sex chromosome. Hereafter, we refer to these ALGs as Merian elements, named after the 17th century lepidopterist and botanical artist, Maria Sibylla Merian (Merian, 1705). Merian elements were named in order of the number of orthologues they carry, ranging from 273 in the largest Merian element (M1) to 19 in the smallest (M31). The sex-linked Merian element, MZ, contains 161 orthologues (table S4).

**Fig. 2:**
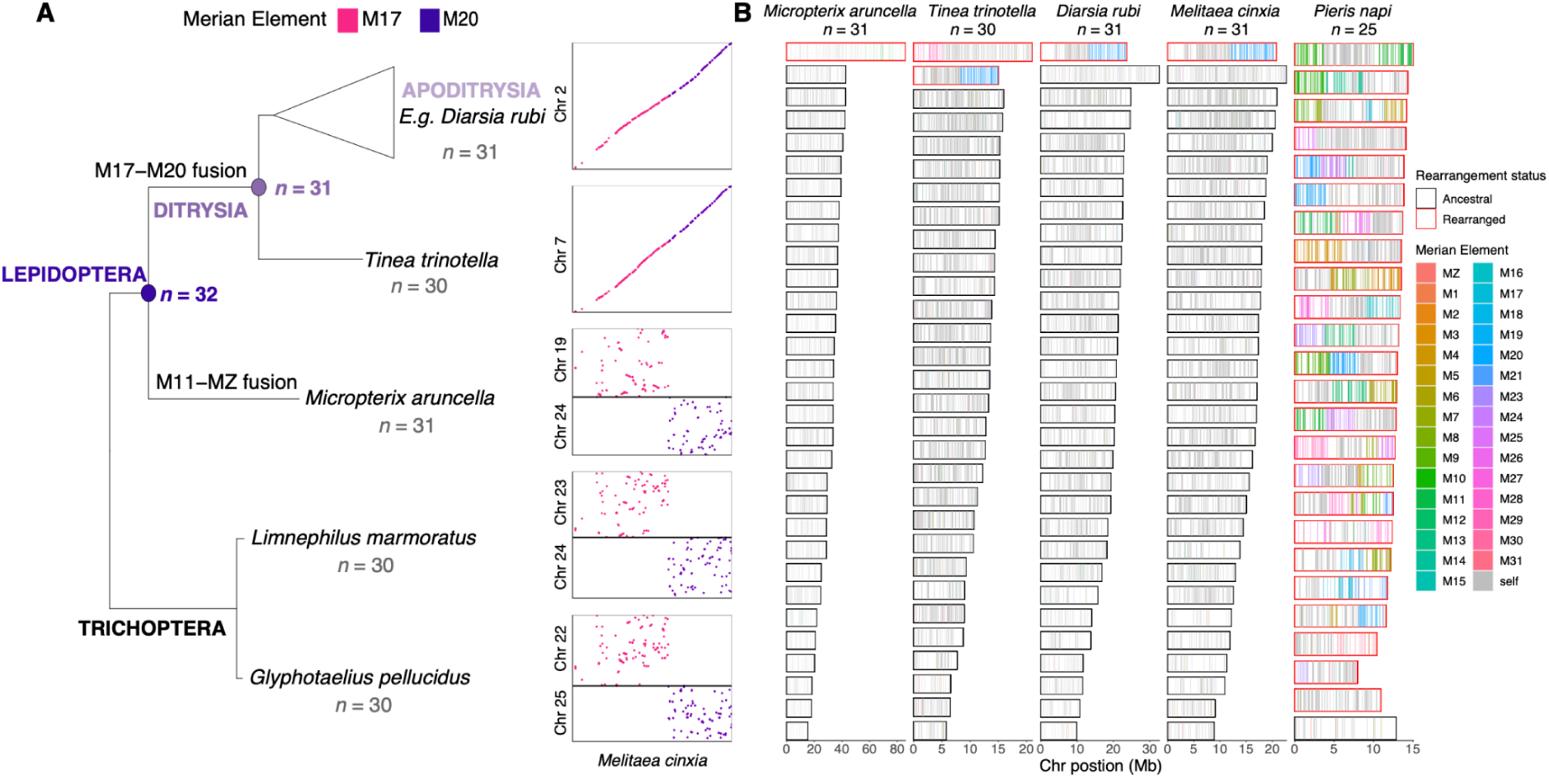
Defining 32 Merian elements. (A) Inferred ancestral karyotype of Lepidoptera and the fusion between M17 and M20 found in all Ditrysia. Phylogeny contains representatives of the two suborders of Trichoptera, *Limnephilus marmoratus* and *Glyphotaelius pellucidus*, in addition to the early-diverging lineage within Lepidoptera, *Micropterix aruncella* and the early-diverging lineage within Ditrysia, *Tinea trinotella* and a representative of *Ditrysia, Diarsia rubi.* To the right of each species in the phylogeny, an Oxford plot of the chromosomes containing orthologues belonging to M17 and M20 in the species is shown relative to *Melitaea cinxia,* which has the chromosome complements of a typical ditrysian species. (C) Merian elements painted across the chromosomes of *Micropterix aruncella, Tinea trinotella, Diarsia rubi, Melitaea cinxia* and *Pieris napi.* Each chromosome is represented by a rectangle within which the position of each ortholog is painted grey if it belongs to the most common Merian element for that chromosome, or else coloured by the alternative Merian element. Chromosomes that have undergone fusions and/or fission events are outlined in red.

An ancient fusion involving M17 and M20 occurred on the branch leading to the last common ancestor of Ditrysia, the most taxonomically and ecologically diverse group of Lepidoptera (Fig. 2A), resulting in the 31 linkage groups observed in the majority of extant ditrysian species. We refer to this fusion as ‘M17+M20’, where the ‘+’ denotes an end-to-end fusion, without mixing of genes. In *Micropterix aruncella*, from the early-branching family Micropterigidae, M17 and M20 are distinct chromosomes and M17 and M20 ALGs were distinct in the last common ancestor of Trichoptera. As the separation of loci defining M17 and M20 is identical in *M. aruncella* and the four Trichoptera, this excludes the possibility that these represent two independent fissions of an ancestral element (Fig. 2A).

We explored the evolutionary dynamics of Merian elements by ‘painting’ the positions of orthologues that define each element onto chromosomes of present-day species (Fig. 2B). With the exception of the ancient M17+M20 fusion that is shared by all ditrysian species, the chromosomes of most species correspond to intact Merian elements. Simple fusion and fission events identified in several species reflected previous cytological karyotype assessments (Robinson, 1971). For example, the chromosomes of *M. aruncella* directly correspond to single, intact Merian elements, with the exception of one Z-autosome fusion (MZ+M11). In *Tinea trinotella*, which have a cytological *n* of 30 (Makino, 1951) we identified a Z-autosome fusion (MZ+M29). Gene-level synteny within each element is highly conserved, even after chromosomal fusion events, including on the ancient M17+M20 (Fig. 2A). More complex rearrangements have occurred in 14 species from 8 lineages. For instance, in *Pieris napi* most chromosomes were made up of segments derived from more than one Merian element and individual Merian elements were fragmented across multiple chromosomes, indicating a history of many fusion and fission events, as proposed previously (Hill *et al*., 2019). In chromosomes that had not undergone rearrangement events, the proportional length of each Merian element was broadly conserved across species (Fig. 3A).

**Fig. 3:**
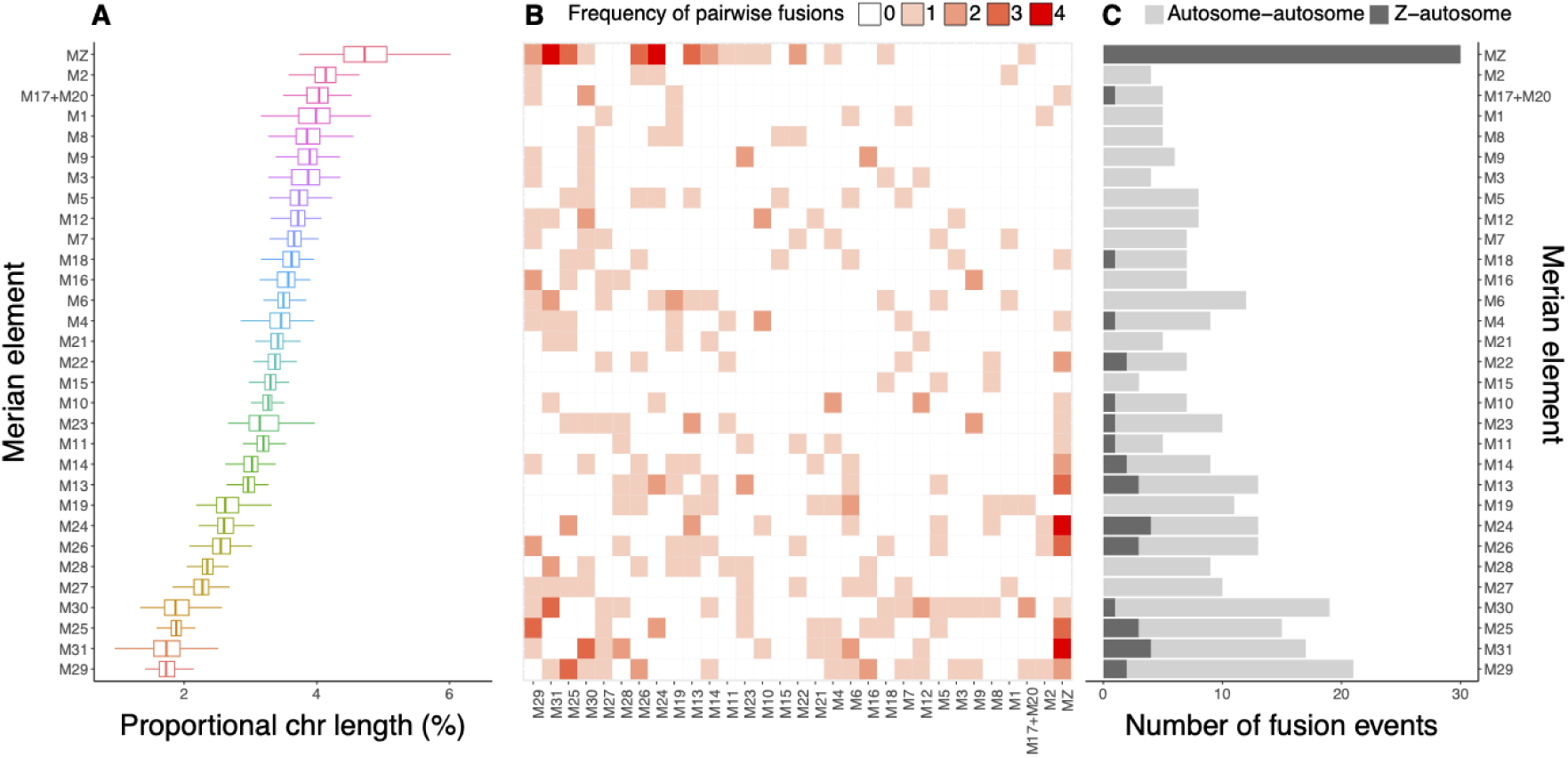
The relationship between Merian element length and tendency to be involved in fusions. (A) Conservation of Merian element length across Lepidoptera. Boxplots of the variation in proportional chromosome length within each Merian element. Only Merian elements that have remained intact (no large-scale rearrangements) are included. Black lines represent standard deviations and black circles represent mean lengths. (B) Matrix of fusion events between pairs of Merian elements, where the shade of red indicates the total number of fusion events per Merian element. (C) Barchart of the number of autosome-autosome and sex chromosome-autosome fusion events that each Merian element is involved in. Merian elements are ordered based on average proportional length across the 210 species.

### The distribution of large-scale rearrangement events across Lepidoptera

Merian elements provide a foundation to infer the dynamics of chromosomal fusions and fissions that generated the karyotypes of extant lepidopteran species and explore pattern and process in chromosome evolution. We inferred the rearrangement histories of 196 species where chromosome painting indicated simple fusions between at most two Merian elements or fission of single Merian elements using two complementary, phylogenetically-aware algorithms that use the co-occurence of orthologues belonging to the same Merian element, which gave the same results.

Excluding the ancient M17+M20 fusion, 54% (106/196 species) have retained intact Merian elements since the last common ancestor of Lepidoptera. We identify 183 simple fusion events and four fission events in the 90 Ditrysian species that deviate from *n*=31 (Fig. 1B). Fission was observed in just three species (*Celastrina argiolus*, *Macaria notata* and *Eupithecia centaureata),* which have one, one, and two fissions, respectively. We also identified a single instance where segments of two Merian elements had fused together, and the remaining portions existed as separate chromosomes, resulting from two fissions (M1 and M6 in *Eupithecia centaureata)* (fig. S7). Most (159, 86%) of the 183 simple fusions appeared to be evolutionarily young, as they were observed in single species. However, 25 fusions mapped to 14 internal nodes were shared by all descendents (Fig. 1A) (table S5,S6). We found that the number of fusions observed in single lineages is significantly greater than expected under a uniform model of evolution. The scarcity of older fusions suggests that lineages with these rearrangements have a reduced probability of persisting over time. We found no instances where reversion of a fusion was a parsimonious explanation of observed karyotypes. We explored whether all Merian elements were equally likely to be involved in fusions. For this analysis, only fusions involving two elements were considered and the ancient fusion observed in all Ditrysia was considered as one unit. We found that some Merian elements are more frequently involved in fusion than other elements (Fig. 3B). The most common fusion pairings were MZ+M31 and MZ+M24 (each with 4 independent occurrences). Strikingly, MZ was involved in the highest number of fusion events (30 independent fusion events). We found that small autosomal elements were involved in a greater number of fusion events than larger ones (Spearman’s rank correlation, *ρ*(29) = −0.6177, *p* = 0.0002136) (Fig. 3C, fig. S8). A bias towards the involvement of smaller chromosomes in fusion events in evolutionary distant lineages has been suggested previously based on analysis of fusions in *B. mori* and *H. melpomene* compared to *M. cinxi*a (Ahola *et al*., 2014). Our analysis suggests that this holds across Lepidoptera and is true for both autosome-autosome fusions and Z-autosome fusions.

### Eight independent lineages have undergone extensive rearrangements

Against the backdrop of strong constraint in karyotype evolution, fourteen species from eight lineages had highly reorganised genomes (Fig 1A). We identified two distinct patterns, exemplified by *Lysandra*, where fission is dominant (Fig. 4A), and by tribe Pierini (Pieridae), where chromosomes have undergone many nested fusion and fission events (Fig. 4B). These species demonstrate that two processes that generate karyotype variation, fission and fusion, can be modulated independently. We found no evidence of polyploidy in any lineage.

**Fig. 4:**
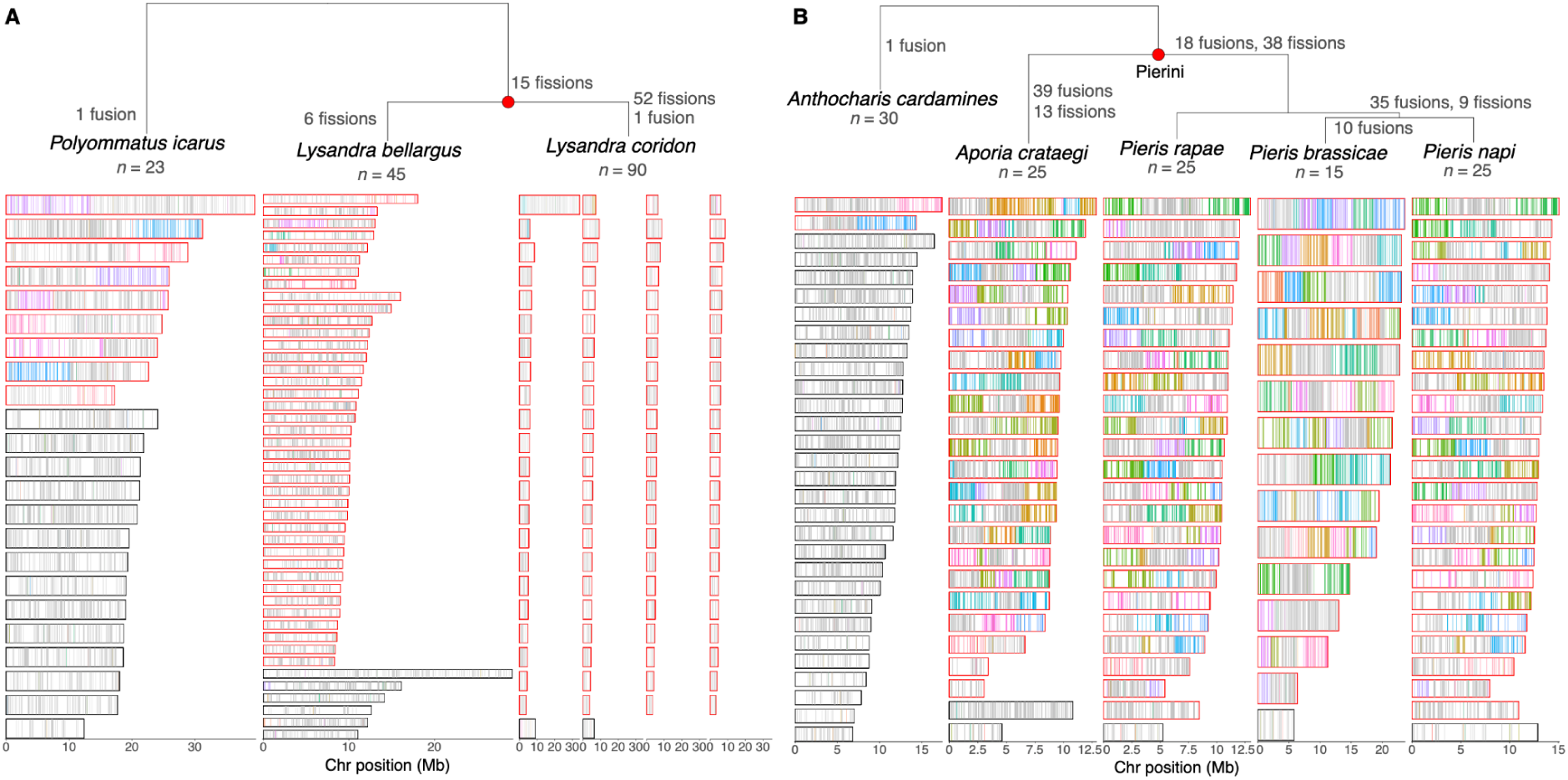
Extensive chromosomal rearrangements in *Lysandra* and Pierini. (A) Relationships of *Lysandra* species with reorganised genomes and sister species *Polyommatus icarus* that has retained intact Merian elements with the exception of 7 fusions shared by all Lycaenids. (B) Relationships of the Pierini species that have reorganised genomes and their sister species, *Anthocharis cardamines,* which is not reorganised. In both panels, Merian elements are painted across the chromosomes of each species. Each chromosome is represented by a rectangle within which the positions of orthologues are painted grey if it belongs to the most common Merian element for that chromosome, or coloured by the alternative Merian element. Chromosomes that have undergone large-scale rearrangements (fusions or fissions) are outlined in red.

To investigate the dynamics of the fission events in *Lysandra,* we reconstructed the events that gave rise to the genome structure of *L. coridon* and *L. bellargus*. Seven pairwise fusions generated a karyotype of *n*=24 in the last common ancestor of family Lycaenidae. Fifteen fissions then generated *n*=39 in the last common ancestor of *Lysandra* (Fig. 4A). Subsequently, *L. bellargus* underwent 6 additional, independent fissions generating *n*=45, and *L. coridon* experienced at least one fission event in 37 of the 39 *Lysandra-*ALGs, resulting in 90 chromosomes. The MZ element did not undergo fission in either species, but fused to a portion of M16 in *L. coridon*. An overwhelming majority of the 90 chromosomes in *L. coridon* map to a single Merian element and show conservation of gene order despite the numerous rearrangement events (fig. S9). The few *L. coridon* chromosomes containing segments from multiple Merian elements derive from fused chromosomes present in the common ancestor of Lycaenidae. A similar pattern of dominance of fission was observed in *Tinea semifulvella*, which has undergone 15 simple fission events, resulting in a karyotype of *n*=45 relative to its sister species, *Tinea trinotella* (*n*=30) (fig. S10).

In Pierini, where chromosomes are mosaics of segments of Merian elements, we inferred a parsimonious set of events that explain the chromosome sets of *Pieris napi, P. rapae, P. brassicae,* and *Aporia crataegi*. A set of fusion and fissions occurred in the last common ancestor of Pierini, and are thus absent in the outgroup *Anthocharis cardamines*. Further fusions and fissions occurred independently in the lineages leading to *A. crataegi* and to the three *Pieris* species (Fig 4B). *P. rapae* and *P. napi* share 25 orthologous, collinear blocks and thus have maintained the same repertoire of fusion and fission events as the last common ancestor of *Pieris* for ∼30 million years (Edger *et al*., 2015). In contrast, *P. brassicae* has undergone 10 further fusion events resulting in a reduced karyotype of *n*=15. With the exception of *P. brassicae*, all *Pieris* species with known karyotypes have haploid chromosome counts between 24 and 27, with most possessing 25 (Robinson, 1971).

Similar to the situation in Pierini; complex, nested rounds of fusion and fission events have shaped the genomes of *Melinaea*. We infer that a series of fusions and fissions occurred in the last common ancestor of *Melinaea*, with further independent fusions and fissions occurring in *M. marsaeus* and *M. menophilus* (fig. S11). Likewise, the genomes of *Brenthis ino* (Nymphalidae) and *Apeira syringaria* (Geometridae) reflect a history of many fusion and fission events, having undergone an estimated total of 33 and 38 events respectively (fig. S12,13). *Leptidea sinapis* (Pieridae) appears to have undergone 29 fusion and 26 fission events, resulting in *n*=48 compared to its close relative, *Anthocharis cardamines*, which has *n*=30 (fig. S14). Two geometrids, *Operophtera brumata* and *Philereme vetulata*, sisters in the dataset, both had highly reorganised genomes. Their last common ancestor had 3 fissions. *O. brumata* then experienced a further 11 fissions and 30 fusions. In contrast, one fusion and 35 fissions occurred in *P. vetulata* (fig. S15). Interestingly, in all highly reorganised lineages, MZ has remained intact with no fissions, and in all lineages but *P. vetulata*, it has fused to one or more autosomal Merian elements.

### Understanding biases in chromosomal fusions in Lepidoptera

Given the preference towards the involvement of smaller Merian elements in fusion events, we asked whether there were compositional differences that vary with chromosome length. We observed striking correlations of GC with chromosome length. In 84% (163/193) of analysed species, we observed a negative correlation between GC content and proportional chromosome length (Spearman’s, p-value < 0.05) (table S8, fig. S16A) with small chromosomes having high GC content. GC content has several drivers, including contributions from repetitive elements, but GC3 (the GC content of the third bases of potentially degenerate codons) is independent of many of these. Only half (48%; 93/184) of the analysed species had higher GC3 values in smaller chromosomes (table S8, fig. S16B) suggesting some of the variation in GC we observed is driven by the density of features such as repeats. Consistent with this, smaller chromosomes have more repeats than large chromosomes (Fig 5A). Negative correlation between chromosome length and repeat density was observed in 93% (180/193) of assayed species (Spearman’s, p-value < 0.05) and ranged in strength from –0.41 in *Notocelia uddmanniana* to ‘0.98 in *Biston betularia* (table S8). High repeat density in smaller chromosomes was not associated with specific repeat types. All major repeat families are enriched in shorter chromosomes, albeit some families more so than others (fig. S17). In contrast to GC content and repeat density, we observed no consistent correlation between coding density and chromosome size (negative correlation was observed in 0.5% (1/184) and negative correlation in 18% (33/184) of species; Spearman’s, p-value < 0.05) (fig. S16C), reflecting previous conflicting trends observed in several Nymphalid species (Martin *et al*., 2019; Shipilina *et al*., 2022).

**Fig. 5:**
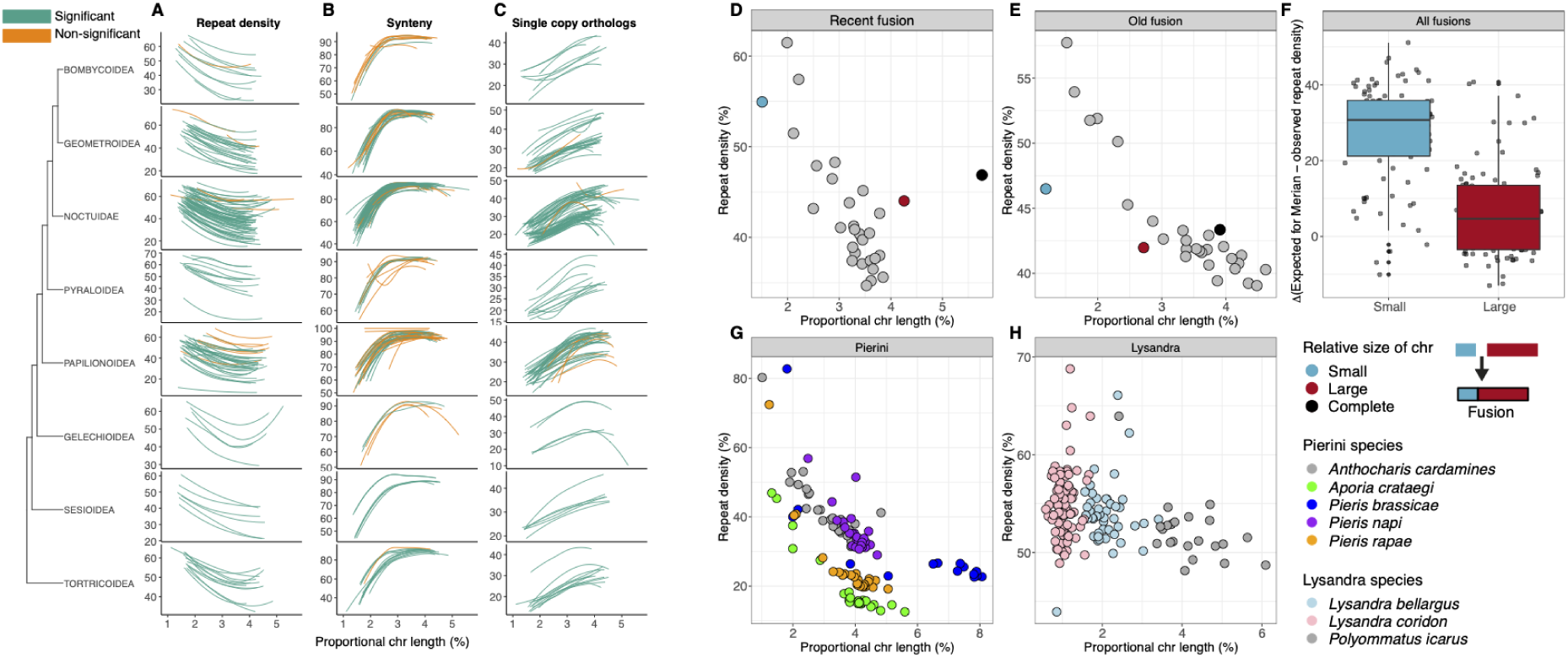
The correlates of chromosome length and sequence features across Lepidoptera. Proportional chromosome length against (A) repetitive element content, (B) synteny defined as the proportion of orthologues which are adjacent in both the reference species *Melitaea cinxia* and the given species, and (C) proportion of chromosomal gene content that is made up of orthologues that are single copy and present across Lepidoptera. The line is coloured green if the correlation with proportional length was significant (Spearman’s rank, p-value < 0.05), or orange if it was non-significant. Only autosomes were included in the correlation analysis. Autosomes were filtered to only retain those that corresponded to intact Merian elements (i.e. had not undergone fusion or fission). Only species with at least 10 autosomes after filtering were analysed and only superfamilies represented by at least 5 species are shown. (D) Proportional chromosome length against repetitive element content for *Agrochola circellaris*, which has a recent fusion, and (E) for *Aphantopus hyperantus*, which has an older fusion. In (F) the difference between the average repeat density of a given Merian element and its current repeat density in the context of a fused chromosome is shown, where small Merian elements are M25, M29, M30, M31. Proportional chromosome length against repetitive sequence content is shown for (G) a set of Pierini species plus the sister species *Anthocharis cardamines* and (H) for species in genus *Lysandra* and the sister species *Polyommatus icarus*.

Smaller chromosomes were also generally less syntenic than longer chromosomes, where synteny is measured as the proportion of adjacent orthologues which are present in both the reference and query species (Fig. 5B). A significant positive correlation (Spearman’s p-value < 0.05) was observed in 68% of species (132/193) with correlation strength ranging from 0.82 in *Limenitis camilla to* 0.37 in *Chrysoteuchia culmella* (Spearman’s, p-value < 0.05) (table S8). Finally, we asked whether the types of genes on small chromosomes are different from those on larger chromosomes. We observed that smaller chromosomes were depleted in single-copy orthologues relative to larger chromosomes in 95% of all analysed species (174/184) (Spearman’s, p-value < 0.05) (Fig. 5C).

In several of these analyses the Z chromosome was an outlier given its relative length (table S9). Unfused MZ had low average GC and GC3 content, in line with GC decreasing with chromosome length (fig. S18 & fig. S19). However, the average repeat content for MZ was higher than expected based on chromosome length alone (fig. S20). While the level of coding density on MZ fell within the range exhibited by autosomes (fig. S21), MZ had a much lower level of synteny than expected based on the scaling of synteny with chromosome length in autosomes (fig. S22). The MZ was also relatively depleted in single copy, conserved genes (fig. S23). Together, these patterns indicate that other evolutionary forces, in addition to the chromosome length, have shaped the content of the Z chromosome.

### Consequences of fusions

The features of Merian elements might be an intrinsic part of their functional biology rather than driven by their relative sizes. Intrinsic function would maintain Merian element-specific feature landscapes in fused chromosomes, while length-related drivers would result in amelioration through time. For phylogenetically recent fusions, the constituent Merian elements had a repeat density similar to that of their ancestral, unfused homologs. For example, in the species-specific M30+M5 fusion in *Agrochola circellaris* we found a repeat-rich M30 segment and a larger and relatively repeat-poor M5 segment. The repeat densities of these segments were in line with the ancestral, unfused expectations (Fig. 5D). Unfused chromosomes had higher repeat densities at their ends, and in recent fusion chromosomes a shoulder of higher repeat density in the area of the fusion, likely a relic from the contributing parts, was evident (fig. S24). *Aphantopus hyperantus* had a phylogenetically older, M29+M14 fusion which was shared by members of subfamily Satyrinae in Nymphalidae. While the M29- and M14-derived segments of the fused chromosome were still distinct in syntenic gene content, they both had repeat densities consistent with the fused chromosome length (Fig. 5E). There was no central shoulder of increased repeat density (fig. S25). In all simple fusions involving one of the four smallest Merian elements, the smaller Merian element tended to have experienced a greater shift in repeat density relative to its unfused ancestor (paired t-test, p-value < 0.01) (Fig. 5F). Thus, the repeat landscape of fused chromosomes evolves over time to reflect that expected of larger chromosomes and patterns of features on chromosomes are largely driven by the relative length of the chromosome and not the identities of the genes carried.

In species with higher chromosome numbers, the average chromosome size will be smaller, and under the model above these will accumulate higher density of repeats. The small, highly reorganised chromosomes of Pierids were indeed repeat-rich relative to the larger chromosomes (Fig. 5G) and the small chromosomes resulting from rampant fission in groups such as *Lysandra* were also repeat-rich (Fig. 5H). Despite the lack of correlation between chromosome number and genome size over all species, accumulation of repeats in species with many, smaller chromosomes was associated with an increase in genome size in *Lysandra* species (fig. S26)*, Leptidea sinapis* (fig. S27), *Philereme vetulata* (fig. S28) and *Tinea semifulvella* (fig. S29). Symmetrically, reduction in chromosome numbers was associated with reduced genome size in Pierini (fig. S30), *Apeira syringaria* (fig. S31) and *Brenthis ino* (fig. S32), but not in *Operophtera brumata* (fig. S28) and *Melinaea* species (fig. S33). It may be that the multiple fusions that reduce chromosome number in these latter species are recent and that insufficient time has passed for repeat content to decrease.

## Discussion

The ongoing revolution in sequencing is enabling major projects such as the Darwin Tree of Life to produce large numbers of chromosomally-complete genomes across eukaryotic diversity (Darwin Tree of Life Project Consortium, 2022; Lewin *et al*., 2022). These rich data permit large-scale, taxon-wide analysis of features and processes previously addressed anecdotally (Blaxter *et al*., 2022). Chromosomal organisation of eukaryotic genomes is one such feature. We analyse over 200 chromosomally-complete genomes and map the evolutionary dynamics of chromosome maintenance, fusion and fission in a holocentric group, the Lepidoptera. We show that the chromosomes of extant species are derived from 32 ALGs, or Merian elements. With the exception of an ancient fusion in the lineage leading to Ditrysia, these ancestral chromosomes have remained intact across most species. These elements have consistent differences in length and genomic features and carry distinct sets of conserved genes which have retained a syntenic order. Merian elements provide a unifying system to explore genomic stasis and change in Lepidoptera, similar to Müller elements of *Drosophila* and Nigon elements of rhabditid nematodes (Muller, 1940; Tandonnet *et al*., 2019; Bracewell *et al*., 2020; Gonzalez de la Rosa *et al*., 2021).

Across Lepidoptera, we find fusions to be sporadic and fissions rarer still. Lepidopteran chromosomes arising from fusions retain syntenic domains reflecting their original element origin. In contrast, holocentric nematodes have a high rate of intrachromosomal rearrangement that leads to rapid mixing of Nigon elements in fused chromosomes (Tandonnet *et al*., 2019; Gonzalez de la Rosa *et al*., 2021). We find smaller Merian elements and the sex chromosome (MZ) are more likely to be involved in fusion events than larger autosomal elements. We have not identified any reversions of fusions back to their unfused homologs, suggesting that reversions are extremely rare. The distinct relative sizes of Merian elements also mean that they evolve differently. In Lepidoptera, each chromosome undergoes one meiotic recombination per generation on average (Yamamoto *et al*., 2008; Davey *et al*., 2017), resulting in smaller Merian elements experiencing higher recombination rates (per base) than longer elements. In addition to reducing linkage disequilibrium and enhancing the efficacy of selection, recombination is mutagenic (Halldorsson *et al*., 2019), and smaller elements will also experience higher mutational pressures. Stability of Merian element size across Lepidoptera means these differences will have had long-term impact on the evolutionary trajectories of the genes and genetic systems each element carries, and that elements that fuse or split will experience a step change in evolutionary rates. Consistent with this, fused Nymphalidae chromosomes have decreased nucleotide diversity compared to their unfused homologues (Cicconardi *et al*., 2021) and raised barriers to introgression (Edelman *et al*., 2019; Martin *et al*., 2019; Mackintosh *et al*., 2023).

In our dataset, MZ was usually longer than autosomal chromosomes and had patterns of repeats, GC proportion and gene content that diverged from expectations derived from the longer autosomes. Because of achiasmatic oogenesis, 67% of the population of MZ but only 50% of the population of autosomal elements undergo recombination each generation. The elevated recombination rate of the Z and haploid exposure in females likely explain the elevated repeat content and GC proportion of MZ compared to autosomal elements (Mongue, Hansen and Walters, 2022). We found that MZ was much more likely to have contributed to fusions than any autosomal element. Sex chromosome-autosome fusions are also overrepresented in rhabditine nematodes (Gonzalez de la Rosa *et al*., 2021), flies (Bracewell *et al*., 2020), vertebrates (Graves, 2016) and plants (Rifkin *et al*., 2021). MZ-autosome fusions may confer selective benefits. Possible drivers are female meiotic drive (Úbeda, Patten and Wild, 2015), sexually antagonistic selection (Charlesworth and Charlesworth, 1980) or deleterious-mutation sharing (Charlesworth and Wall, 1999; Antonovics and Abrams, 2004; Jay *et al*., 2022). The set of thirty independent MZ-autosome fusions described here presents a valuable dataset to dissect the drivers of the rate of molecular evolution in sex chromosomes and, for fusions, illuminate the forces that shape autosomes. The resistance of MZ to fission in species where fission is dominant also requires deeper exploration.

Why have Merian elements remained largely stable in content and gene order through ∼250 million years (Kumar *et al*., 2017; Kawahara *et al*., 2019) of lepidopteran evolution? Species with holocentric chromosomes are theoretically more permissive to karyotypic change. This is reflected in some holocentric groups, such as *Carex* sedges where karyotype evolution is rapid (Hipp, 2007; Roalson, 2008), but is not seen in insects where systematic differences between fusions and fission rates in holocentric versus monocentric groups are not found (Ruckman *et al*., 2020). One potential constraint on the fixation of rearrangements is the ability to undergo meiosis. Individuals heterozygous for rearrangements can be sterile due to unbalanced segregation (heterozygote disadvantage), leading to underdominance (Grant, 1981; King, 1995; Vershinina and Lukhtanov, 2017). Structural heterozygosity impacts reproductive fitness in holocentric *Caenorhabditis elegans* nematodes and *Carex* (Dernburg, 2001; Escudero *et al*., 2016). Pairing centre activity has been hypothesised to impose a constraint on karyotype evolution (MacQueen *et al*., 2005; Rog and Dernburg, 2013; Carlton, Davis and Ahmed, 2022). In *C. elegans* the two classical roles of centromeres, as sites of kinetochore assembly and of homologue pairing, have been separated. Kinetochores assemble in broad domains along *C. elegans* chromosomes in areas of low transcriptional activity, but homologue pairing is restricted to discrete regions, meiotic pairing centres, enriched for short sequence motifs. While pairing centres have not been defined in Lepidoptera, investigation of *Bombyx mori* identified similarity to *C. elegans*, with non-sequence-specific kinetochore assembly in regions with low transcriptional activity (Senaratne *et al*., 2021). Understanding lepidopteran kinetochore and pairing centre biology will illuminate the roles of these basic systems in constraining or promoting chromosome number evolution.

Merian elements may be maintained because the genes they carry need to be colocated. Genes in close linkage for long evolutionary periods may evolve *cis*-regulation which would be disrupted by large-scale rearrangement (Peng, Pevzner and Tesler, 2006). It has been suggested that the fragments of Merian elements that fused to produce the highly rearranged chromosomes of *Pieris* species represent sets of genes with related functions, and these gene networks present a constraint (Hill *et al*., 2019). Consistent with this, fusions disrupted patterns of inter- and intra-chromosomal contacts in mouse germ cells (Vara *et al*., 2021) and rearrangement hotspots exist at the boundaries of topologically-associated domains in mammalian chromosomes (Damas *et al*., 2022) with deletion of these domains disrupting development in mice (Rajderkar *et al*., 2023). However topologically-associated domains are usually much shorter than individual chromosomes, and so are unlikely to offer a complete explanation of Merian elements conservation.

Chromosomal evolution in Lepidoptera is not homogenous: in a background of stasis are lineages that have experienced major change. As chromosome number evolution positively correlates with speciation rate in Lepidoptera (de Vos *et al*., 2020), variation in processes that influence the rate of karyotypic change will impact the rate of speciation. For most species, chromosome number has been constrained to *n*=32-31, suggesting a genetic counting mechanism. For the eight identified lineages with major changes in karyotype, this constraint has been lifted. We classify these lineages into: (i) extensive fission of autosomal elements resulting in many small autosomes and a large, intact MZ, or (ii) a burst of fission and fusion events. In the latter, we also identify re-establishment of the normalising counter, albeit at different values of *n*. For example, after initial fission and fusions, *Pieris* species restabilised at *n*=25. There are thus three processes which generate lepidopteran chromosomal complements: a normalising *counter* that maintains the status quo, a *scission* mechanism that splits elements and a *fusion* process that promotes joining of elements and fragments. These correspond to the normative, ascending and descending disploidy parameters of established karyotype evolution models (Mayrose and Lysak, 2021). We find that these processes can be separately modified in different species and lineages, with the counter dominant and fission and fusion largely repressed in most species, and, for example, scission derepressed in *Lysandra*, and scission and fusion derepressed but fusion more recently dominant in *P. brassicae*. These processes have biases related to Merian element size and identity. Smaller chromosomes are more likely to fuse even when the counter is dominant, and the MZ and autosomes fused to it are insulated from fission.

Elevated rates of fixation of rearrangements may be a product of neutral processes such as genetic drift of mildly deleterious and/or underdominant changes during sustained periods of low effective population size (Templeton, 1981). Functional differences in core chromosome biology could also drive change. In parrots (Aves; Psittaciformes), frequent rearrangements have been linked to the loss of genes involved in the repair of double-strand breaks and maintenance of genome stability (Huang *et al*., 2022). The existence of lepidopteran lineages where fission and fusion rates have been individually modified will permit detailed investigation of their mechanistic bases. We note that several species with highly reorganised genomes display variable karyotypes between populations (Coutsis, De Prins and De Prins, 2001; Lukhtanov *et al*., 2011). Mating between individuals with highly divergent karyotypes can produce fertile offspring, suggesting that meiosis in some lepidopterans can tolerate heterozygosity for multiple rearrangements (Kitahara, 2008, 2012; Lukhtanov *et al*., 2018). However the persistence of hybrid zones between demes with different karyotypes suggests that there is a fitness cost in hybrids (Lukhtanov *et al*., 2011). Transposable elements are hypothesised to facilitate high rates of chromosome fusion (Ahola *et al*., 2014; Mathers *et al*., 2021; Lohse *et al*., 2022) by promoting deletion, translocation, and inversion (Miller and Capy, 2004). The smaller lepidopteran autosomes, more frequently involved in fusions, do have higher repeat content, but MZ, which has relatively low repeat density and fuses frequently, does not. An enrichment of LINEs at fusion boundaries observed in *L. sinapis* (Höök *et al*., 2023) may be an unameliorated relic of the fusion. Moreover, analysis of the *P. napi* genome found no enrichment of repeats at fusion boundaries and no repeat class was expanded compared to other species (Hill *et al*., 2019).

While the impacts of karyotype on evolutionary trajectories may be indirect, their effects can be profound. All other things being equal, change in karyotype between species is unlikely to be neutral. Fundamentally, change likely promotes speciation (de Vos *et al*., 2020). However, the pattern of overall stasis indicates that lineages with highly variant karyotypes may be at a macroevolutionary disadvantage despite any short-term speciation advantage. We highlight that higher chromosome counts mean more recombination and thus potentially faster evolutionary rates (or more effective selection) overall. This effect will be particularly marked for genes on the elements directly involved in fusions and fissions, and genome-wide in species with widespread changes such as the eight lineages with extensive rearrangement. By analysing 210 chromosomally-contiguous genomes across Lepidoptera, we provide a comprehensive characterisation of genome structure, facilitating deeper investigation into the drivers of rearrangements and lay the foundations for large-scale investigations of genome evolution across the tree of life.

## Materials and methods

### Chromosomal genome assemblies, annotations and transposable elements identification

We downloaded all representative chromosome-level reference genomes for Lepidoptera and Trichoptera that were available on INSDC on 27th June 2022. Of these 212 lepidopteran genomes and 4 trichopteran genomes, 191 were generated by the Darwin Tree of Life Project (Darwin Tree of Life Project Consortium, 2022). Accession numbers and references for all genomes are given in table S1. For species generated by the Darwin Tree of Life project that do not have a reference, the methods were the same as for (Boyes *et al*., 2021). We used the primary assembly for all analyses. The speciose Noctuoidea (71 species) and the intensely studied Papilionoidea (51 species) contribute most to the genomes.

Gene annotations were generated by Ensembl (Cunningham *et al*., 2022) (http://rapid.ensembl.org) for 201 species (table S3). Species that had publicly available RNA sequencing (RNA-Seq) data were annotated using Genebuild, which makes use of both RNA-seq and protein homology evidence. For species that did not have transcriptomic data, the genomes were annotated using BRAKER2 (Brůna *et al*., 2021) using protein homology information as evidence. Protein data consisted of OrthoDB (v11) data (Zdobnov *et al*., 2021) for Lepidoptera combined with all lepidopteran proteins with protein evidence levels 1 or 2 from UniProt (UniProt Consortium, 2019) (where level 1 or 2 represent evidence from either proteomic or transcriptomic data). Details of each annotation are provided in table S3. The gene sets contained between 9,267 (*Tinea trinotella*) and 23,879 *(Miltochrista miniata*) protein-coding genes and between 15,416 (*Erynnis tages*) and 41,125 (*Dendrolimus puncatus*) transcripts. Transposable elements (TE) were identified using the earlGrey TE annotation pipeline (v1.2) (Baril, Imrie and Hayward, 2021) on each genome as described in (Baril and Hayward, 2022), with the Arthropoda library from Dfam release 2.5 (Jurka *et al*., 2005; Hubley *et al*., 2016). Repeat annotations are provided in Supplementary Data 2.

Two genomes were excluded from further analysis due to quality issues. The first, *Zerene cesonia* (GCA 012273895.2), contained 246 unlocalised scaffolds that contained 351 BUSCOs. The high number of BUSCOs in these scaffolds means that erroneous rearrangement events would be inferred if this genome were to be included. In the second, *Cnaphalocrocis medinalis* (GCA 014851415.1), the majority of genes belonging to the M30 Merian element were present on unlocalised scaffolds. We identified two additional genomes that contained minor misassembly issues that we were able to address prior to downstream analysis. In *Dendrolimus kikuchii* (GCA 019925095.1), we found three scaffolds with a high proportion of duplicated BUSCOs (the majority of which corresponded to the M30 Merian element), indicating that they represented haplotypic duplication. When we removed these scaffolds from the assembly, we successfully recovered a fusion between M30 and MZ that would have otherwise been missed. In *Spodoptera frugiperda* (GCA 011064685.2), we removed an unlocalised scaffold that contained 22 BUSCOs prior to downstream analyses to avoid inferring a fission event in this species due to assembly issues.

### Phylogenetic tree reconstruction

We used BUSCO v.5.4.3 (using the metaeuk mode and the lepidoptera odb 10 dataset) (Simão *et al*., 2015) to identify single-copy orthologues in each genome. We used busco2fasta.py (available at (https://github.com/lstevens17/busco2fasta) to identify 5,046 BUSCO genes that were single-copy and present in at least 90% of the genomes. We aligned the protein sequences of these BUSCOs using MAFFT v7.475 (Katoh and Standley, 2013) and trimmed alignments using trimal v1.4 (Capella-Gutiérrez, Silla-Martínez and Gabaldón, 2009)) with parameters -gt 0.8, -st 0.001, -resoverlap 0.75, -seqoverlap 80. A total of 4,947 alignments passed the alignment thresholds. We concatenated the trimmed alignments to form a supermatrix using catfasta2phyml (available at https://github.com/nylander/catfasta2phyml). We provided this supermatrix to IQ-TREE v2.03 (Nguyen *et al*., 2015) to infer the species tree under the LG substitution model (Le and Gascuel, 2008) with gamma-distributed rate variation among sites and 1000 ultrafast bootstrap replicates (Hoang *et al*., 2018). The tree was rooted on the node separating Trichoptera and Lepidoptera and visualised alongside genome size and chromosome number information using *ggtree* v3.0.2 (Yu *et al*., 2017, 2018). The phylogeny is provided in Supplementary Data S2.

To test for a correlation between genome size and chromosome number, we employed a phylogenetic linear model using the R package *phylolm* v2.6.2 (Ho and Ané, 2014) with genome size as the response variable and chromosome number as a fixed factor. To account for shared ancestry between species, the phylogenetic tree described above was included. The most appropriate model for the error terms was identified as Ornstein-Uhlenbeck (OU) by fitting all implemented models that allow for measurement error and then selecting the best-fitting model via the AIC values.

### Defining and visualising Merian elements

We inferred the ancestral lepidopteran linkage groups using syngraph (https://github.com/DRL/syngraph) (using a threshold of 5 orthologues and using the mode that infers fusions and fission events) using the BUSCO-derived single-copy orthologues and the phylogeny derived from all 210 chromosomal lepidopteran genomes and 4 chromosomal trichopteran genomes. As described in (Mackintosh *et al*., 2023), syngraph uses parsimony to infer the arrangement of orthologues in the last common ancestor of species triplets. Syngraph works from the tips towards the root to infer ancestral linkage groups (and fusion and fission events, discussed below) at each internal node in the tree. We used the ancestral linkage groups inferred by syngraph in the last common ancestor of all Lepidoptera in our analysis, which we termed Merian elements. We named Merian elements in ascending order based on the number of orthologues contained (M1 - M31). The group of orthologues that represented the ancestral Z chromosome were named MZ. We “painted” the chromosomes of each extant species to show the distribution of these Merian elements using custom scripts (https://github.com/charlottewright/lep_busco_painter). Merian paints for each species are provided in Supplementary Data S3. We also visualised synteny between pairs of species using Oxford plots generated using custom scripts (available at: https://github.com/charlottewright/Chromosome_evolution_Lepidoptera_MS).

### Inferring fusion and fission events

We inferred simple fusion and fission events (defined as those that involve complete Merians and did not appear to be nested) using two complementary approaches: syngraph and lep_fusion_fission_finder (LFFF) https://github.com/charlottewright/lep_fusion_fission_finder. As discussed above, syngraph infers ancestral linkage groups at each internal node in the tree along with any fusion and fission events that occurred at each branch. In contrast, LFFF uses a set of ancestral linkage groups (in this case, the Merian elements inferred by syngraph) to identify fused or split chromosomes in extant species only. To do this, LFFF identifies the most common Merian element in non-overlapping windows of a given size. Fused chromosomes are identified as those containing windows assigned to two or more Merians (and the position along the chromosome where Merian-element identity switches is recorded as the fusion position). Split chromosomes are identified as those where a Merian is assigned to two or more unfused chromosomes. Fusion and fission events are then inferred by mapping these fused and split chromosomes onto the phylogeny. We identified the optimal number of orthologues as a threshold in both syngraph and LFFF by manually assessing the inferred events. At low thresholds (<17), small rearrangement events or unlocalised scaffolds are often identified as fused chromosomes or split chromosomes. However, at higher thresholds (>17), small or split Merians are often erroneously excluded. We identified the optimal threshold at 17 for both syngraph and LFFF. Using this threshold, we obtained nearly identical results with both approaches, with the only differences being due to how fusions involving more than two Merians are denoted (table S5, table S6). The genomes of species which had one or more examples where orthologues belonging to a single Merian element were present along more than one chromosome, and where such chromosomes are not the product of simple fission events, were classified as highly-rearranged species with complex rearrangements, and so were analysed separately. Similarly, species with genomes resulting from many fission events, leading to at least one chromosome with fewer Merian-defining orthologues than our threshold (<17), were classified as highly-rearranged and so analysed separately. To analyse these species, Syngraph was run on the complete set of 210 lepidopterans using a lower, more sensitive threshold of 5 orthologues.

We tested whether an excess of fusions was inferred to be specific-specific, i.e. occurred along external branches, by simulating a null distribution of fusion events over the lepidopteran phylogeny using a custom script (available at: https://github.com/charlottewright/Chromosome_evolution_Lepidoptera_MS). To do this, the branch lengths were recorded over the phylogeny and whether the branch was external or internal. Then, a random sample of 183 fusions were weighted by branch lengths, with the assumption that fusions happen uniformly across the tree. The number of the 183 fusions that were on external branches versus internal branches was then recorded and compared to the observed number of events on external branches.

The strength of rank-based correlation between proportional chromosome length and frequency of fusion events was calculated using Spearman’s rank implemented in the R package *stats* (v.4.1.0), with a p-value < 0.05 cutoff to assess significance.

### Describing feature distributions across chromosomes

We calculated the distribution of sequence features (GC, repeat density and coding density) along each chromosome 100 kb windows. GC content per 100 kb was calculated using fasta_windows (https://github.com/tolkit/fasta_windows). For other features, a BED file specifying the start and end of each 100 kb window was generated for each genome with BEDtools v2.30.0 (Quinlan and Hall, 2010). Repeat density was calculated using BEDtools coverage and the repeat annotation file produced by earlGrey. To calculate coding density, we filtered the GFF3 files using AGAT v1.0.0 (Dainat *et al*., 2022) to retain only the longest transcript per gene. As a quality check, we excluded CDS sequences that were not divisible by three using a custom Python script (available at https://github.com/charlottewright/genomics_tools). The resulting filtered GFF3 files were used with BEDtools coverage to calculate CDS density in 100 kb windows.

We also calculated the density of each feature by splitting each chromosome into 100 windows. First, a BED file specifying the position of each window along chromosomes was made using BEDtools makewindows with the fasta index file generated from samtools index (v1.7) (Li *et al*., 2009). Repeat density was then calculated per window using BEDtools coverage (v2.30.0) (Quinlan and Hall, 2010). GC per window was calculated from the output from running fasta_windows on 100kb windows, using a custom Python script (https://github.com/charlottewright/Chromosome_evolution_Lepidoptera_MS) and the BED file containing the positions of each window.

### Describing feature distributions between chromosomes

We calculated the average density of various features (GC, GC3, repeat density, coding density, synteny, and proportion of single-copy orthologues) in each chromosome (table S7).

The average GC content of each chromosome was calculated using fasta_windows https://github.com/tolkit/fasta_windows. To calculate the average GC3 value per chromosome, the GC3 value for each coding sequence was calculated using gff-stats (https://github.com/charlottewright/gff-stats/) and these values were used to calculate the average per chromosome using a custom Python script (available at https://github.com/charlottewright/genomics_tools/). Average repeat density per chromosome was calculated using BEDtools (v2.30.0) (Quinlan and Hall, 2010).

We calculated the degree of synteny, defined as conserved gene order, per chromosome using a custom Python script (available at https://github.com/charlottewright/genomics_tools/). We calculated synteny as the proportion of adjacent gene pairs that have collinear orthologues in a corresponding species. We used the BUSCO genes defined previously and calculated synteny in each species relative to *Melitaea cinxia*.

The proportion of conserved single-copy orthologues relative to multi-copy orthologues and species- or clade-specific genes was inferred from the annotated proteins obtained from Ensembl. We first filtered the GFF3 files for each species using AGAT to contain only the longest isoform per protein-coding gene. We filtered the corresponding protein files using fastaqual_select.pl (https://github.com/sujaikumar/assemblage). We then clustered all protein files into orthologous groups using OrthoFinder v2.5.4 (Emms and Kelly, 2019). By analysing these groups, we found that the annotation for one species, *Pieris napi*, was missing many orthologues that were present in the vast majority of the other annotations (fig. S2). We, therefore removed this annotation from the dataset and re-inferred orthologues with Orthofinder. We identified 4,946 orthologous groups that were duplicated or missing in no more than 10% of species. We then classified each gene as either single-copy multi-copy or clade-specific using a custom Python script (available at https://github.com/charlottewright/genomics_tools/). The classified genes were then used to calculate the proportion of genes per chromosome that were classified as single-copy versus non-single copy using a custom Python script (available at https://github.com/charlottewright/genomics_tools/)

To compare the density of these features across lepidopteran chromosomes, we considered only those species that contained 10 or more chromosomes that had not undergone a fusion or fission event (which left 193 species). Nine of these species did not have publicly available gene annotation and so coding density, GC3 and proportion of single-copy orthologues could not be analysed. For each feature, the strength of the correlation between the feature value and proportional chromosome length (calculated as the chromosome length divided by the genome size) using the cor.test function in R using Spearman’s rank correlation and a cut-off of p-value < 0.05 was used to assess significance (Best and Roberts, 1975). The strength of rank-based correlation between chromosome length and each feature density was calculated using Spearman’s rank implemented in the R package *stats* (v.4.1.0), with a p-value < 0.05 cutoff to assess significance.

### Repeat analysis within fusion chromosomes

To understand the effect of fusion on the repeat content of fused chromosomes, we chose fusions that involved M31, M30, M29 or M25 (which are the Merians with the lowest proportional length and were therefore expected to contain the highest repeat content). We expected the chromosomes involved in these fusion events to display the largest difference in repeat content prior to the fusion event. We created a BEDfile for each fused chromosome containing two windows, split at the fusion points that were defined by LFFF previously. The average repeat content for each window was calculated using BEDtools coverage. The difference between the repeat content of the larger in length Merian element and the smaller Merian element was statistically compared with a paired t-test as implemented in the R package stats v4.1.

## Acknowledgements

Authors from the Wellcome Sanger Institute were supported by Wellcome Trust award 220540/Z/20/A, ‘Wellcome Sanger Institute Quinquennial Review 2021-2026,. For the purpose of Open Access, the authors have applied a CC BY public copyright licence to any Author Accepted Manuscript version arising from this submission. We acknowledge the many people and groups that generated the publicly-available chromosomal genomes that were used in this work, including the Darwin Tree of Life project coordinated by the Wellcome Sanger Institute (Darwin Tree of Life Project Consortium, 2022). We thank Matthieu Moffato for help with running software at scale and are grateful to Tree of Life colleagues for reading and commenting on an earlier draft of this manuscript.

