## Supplementary_materials for "Chromosome evolution in Lepidoptera"

**The PDF file includes:**

Supplementary Text
Fig. S1 to S29
Tables S1 to S9
References

**Other Supplementary Material for this manuscript includes the following:**

Tables S1 to S9

### Table of Contents


**Supplementary Text** 3

**Supplementary Tables** 5

Table S1 (separate file). Genome assemblies information. 5
Table S2 (separate file). W chromosome information 5
Table S3 (separate file). Gene annotation information 5
Table S4 (separate file). Assignment of 4,112 single copy orthologs to Merian elements 5
Table S5 (separate file). Inferred fusion and fission events with Syngraph 5
Table S6 (separate file). Inferred fusion and fission events with LFSF 5
Table S7 (separate file). Feature statistics for all 6,254 chromosomes in the dataset. 5
Table S8 (separate file). Genomic correlates of chromosome length across 193 species. 5

**Supplementary Figures** 7

Fig. S1: Assessing quality of gene annotations using the number of missing conserved single-copy genes 7
Fig. S2: Assessing quality of gene annotations using the number of duplicated conserved single-copy genes 8
Fig. S3: Distribution of haploid chromosome number in Lepidoptera from 210 species 9
Fig. S4: Distribution of genome size (Mb) in Lepidoptera from 210 species 10
Fig. S5: Genome size is not correlated with haploid chromosome number 11
Fig. S6: Sequence patterns in Lepidopteran chromosomes. 12
Fig. S7: Merian elements painted across the chromosomes of *Eupithecia centaureata* demonstrate fission and fusion involving M1 and M6. 13
Fig. S8: Relationship between proportional length and frequency of fusion events 14
Fig. S9: Conservation of gene order in *Lysandra coridon* relative to *Polyommatus icarus* despite numerous fission events. 15
Fig. S10: Merian elements in two Tinea species relative to *Micropterix aruncella* 16
Fig. S11: Merian elements in two Melinaeae species relative to *Danaus plexippus* 17
Fig. S12: Merian elements in *Brenthis ino* relative to *Fabriciana adippe* and *Boloria selene*. 18
Fig. S13: Merian elements in *Apeira syringaria* relative to *Selenia dentaria* 19
Fig. S14: Merian elements in *Leptidea sinapis* relative to *Anthocharis cardamines* 20
Fig. S15: Merian elements in *Operophtera brumata* and *Philereme vetulata* relative to Hydriomena furcata 21
Fig. S16: Correlation of GC, GC3 and coding density with chromosome length. 22
Fig. S17: Relationship between repetitive element density and proportional chromosome length for each major class of transposable elements 23
Fig S18: Negative relationship between GC and proportional chromosome length of each Merian element 24
Fig S19: Negative relationship between GC3 and proportional chromosome length of each Merian element 25
Fig S20: Negative relationship between repeat density and proportional chromosome length of each Merian element 26
Fig S21: Variation in the coding density against proportional chromosome length of each Merian element 27
Fig S22: Positive relationship between the level of synteny and proportional chromosome length of each Merian element 28
Fig S23: Positive relationship between the proportion of single copy orthologs and proportional chromosome length of each Merian element 29
Fig S24: Repeat density across fused chromosomes and their unfused homologs in pairs of sister species. 30
Fig S25: Repeat density across the fused chromosome in *Aphantopus hyperantus* 31
Fig S26: Relationship between repeat density, genome size and chromosome size in Lysandra 32
Fig S27: Relationship between repeat density, genome size and chromosome size in Leptidea 33

Fig S28: Relationship between repeat density, genome size and chromosome size in *Philereme vetulara* and *Operophtera brumata* 34
Fig S29: Relationship between repeat density, genome size and chromosome size in Tinea 35
Fig S30: Relationship between repeat density, genome size and chromosome size in Pierini 36
Fig S31: Relationship between repeat density, genome size and chromosome size in Apeira 37
Fig S32: Relationship between repeat density, genome size and chromosome size in Brenthis 38
Fig S33: Relationship between repeat density, genome size and chromosome size in Melinaea 39
**References 40**

#### Supplementary Text

Section 1: Dataset selection

The genomes used in this study were generated from a mix of female (heterogametic, ZW) and male (homogametic, ZZ) specimens. Therefore, W chromosomes are not present in all assemblies. Sixty one assemblies were generated from females, of which 27 assemblies contained a single scaffold assigned to a W, with mean size of 14.57 Mb (table S2). The remaining assemblies contained 2-84 scaffolds corresponding to the W chromosome. Two assemblies were inferred to be Z0 due to the absence of any W-linked sequence. As the W chromosome is largely composed of repetitive sequence, it was not included in analyses of chromosome structure.

Section 1: Identification of transposable elements

Transposable elements identified with EarlGrey (Baril, Imrie, and Hayward 2022) occupy between 67.5% (*Micropterix aruncella*) and 7.4% *(Danaus plexippus*) of each genome with an average of 41.7% (SD=10.8%) (table S8). As previously described for several species, we find that LINEs are the most prevalent transposable element in lepidopteran genomes, followed by rolling circle elements (table S9) (Baril and Hayward 2022; Mackintosh et al. 2022). Most species show a higher abundance of DNA elements than LTRs and SINEs but SINES are enriched in the genomes of some species.

Section 1: Phylogeny

The dataset used for the phylogeny included a scaffold-level Trichopteran genome (*Hydropsyche tenuis*) in order to increase the taxonomic breadth of Trichoptera used as an outgroup to Lepidoptera. The phylogeny is consistent with previously published time-calibrated phylogenies, including a recent, comprehensive molecular analysis of lepidopteran phylogeny (Kawahara et al. 2019). All 30 families were recovered as monophyletic and *Micropterix aruncella* (Superfamily: Micropterigidae) was positioned as the earliest diverging within Lepidoptera as expected (Kawahara et al. 2019).

Section 2: Assignment of orthologs to Merian elements

The assignment of the remaining orthologs to a Merian element or as absent in the last common ancestor will likely be possible with the sequencing of further early-diverging relatives including species of Micropterigoidea, Agathiphagoidea, and Heterobathmioidea. Given that all species that were used to build the lepidoptera odb10 set are from Dytrisia, it is likely that many were simply not present in the last common ancestor of Lepidoptera.

Section 3: Filtering the dataset when inferring fusion and fission events

Before inferring fusion and fission events, the distribution of Merian elements were visualised across the chromosomes of all 210 lepidopteran genomes in order to check for any data quality issues due to misassembly. This enabled us to be confident that our resulting inferred fusion and fission events were not artefacts due to misassembly. Most genomes were high quality, with only chromosomal scaffolds containing multiple single copy orthologs as expected. However, three genomes contained unlocalised scaffolds with multiple single copy orthologs which belonged to Merian elements. First, *Spodoptera frugiperda* contained a scaffold (WMCG01000038.1) which contained a set of orthologs corresponding to Merian element 5. To prevent this scaffold from being called as a fission event, it was manually removed from the assembly for subsequent analyses.

The second, *Dendrolimus kikuchii,* contained two scaffolds (JAHHIN010000030.1 and JAHHIN010000032.1) which contained 18 and 7 BUSCOs respectively, of which 16 and 6 BUSCOs were duplicated. This suggested that these scaffolds are the result of unpurged heterozygosity. As their presence prevented a fusion between Merian Z and Merian 31 from being inferred these two scaffolds were manually removed from the assembly for subsequent analyses.

#### Supplementary Tables

**Table S1 (separate file). Genome assemblies information.**

Table contains the assembly name, species, data source, GCA accession number, assembly level, scaffold N50 and BUSCO completeness (lepidoptera odb10). Also contains genome size, chromosome number and taxonomic information.

**Table S2 (separate file). W chromosome information**

Table contains the assembly name, observed sex from the sample sequenced, sex chromosomes identified in the assembly, number of W contigs and size of the W for assemblies where a single contig was assigned as the W.

**Table S3 (separate file). Gene annotation information**

Table specifies whether each annotation was generated using BRAKER2 or Genebuild. The generation date of each annotation is also listed, to provide version information.

**Table S4 (separate file). Assignment of 4,112 single copy orthologs to Merian elements**

**Table S5 (separate file). Inferred fusion and fission events with Syngraph**

**Table S6 (separate file). Inferred fusion and fission events with LFSF**

**Table S7 (separate file). Feature statistics for all 6,254 chromosomes in the dataset.**

Table includes species name, assigned Merian elements, rearrangement status, GC3, synteny, coding density, repeat density, chromosome length and genome size.

**Table S8 (separate file). Genomic correlates of chromosome length across 193 species.**
Strength of correlation between coding density/ repeat density/ GC3/ synteny/ single copy orthologs versus proportional chromosome length for all analysed lepidopteran species.

**Table S9 (separate file). Mean feature per Merian element across the dataset**
Table specifies the mean feature density and standard deviation for a set of features - scaled GC content, scaled GC3 content, scaled repeat density, scaled coding density, synteny and proportion of single copy, conserved orthologs relative to multi copy and singleton genes - per Merian element. Only chromosomes corresponding to intact Merian elements (i.e. have not undergone fusion or fission) in the 210 lepidopteran genomes were included in these statistics.

##### Full list of links to raw data and source code

Reference genomes:
https://www.ncbi.nlm.nih.gov/ - GCA accession numbers are given in table S1.

Gene annotations:
Rapid.ensembl.org - versions listed in table S3

Supplementary data sets (Wright 2023):
https://zenodo.org/record/7925505
Supplementary Data S1: Repeat annotations for each genome analysed
Supplementary Data S2: Phylogenetic tree for the analysed genomes
Supplementary Data S1: Merian elements painted across the chromosomes of each analysed species

Script and methods used to analyse fusion chromosomes:
https://github.com/charlottewright/lep_fusion_fission_finder
https://github.com/charlottewright/lep_busco_painter
https://github.com/charlottewright/Chromosome_evolution_Lepidoptera
https://github.com/charlottewright/genomics_tools

#### Supplementary Figures

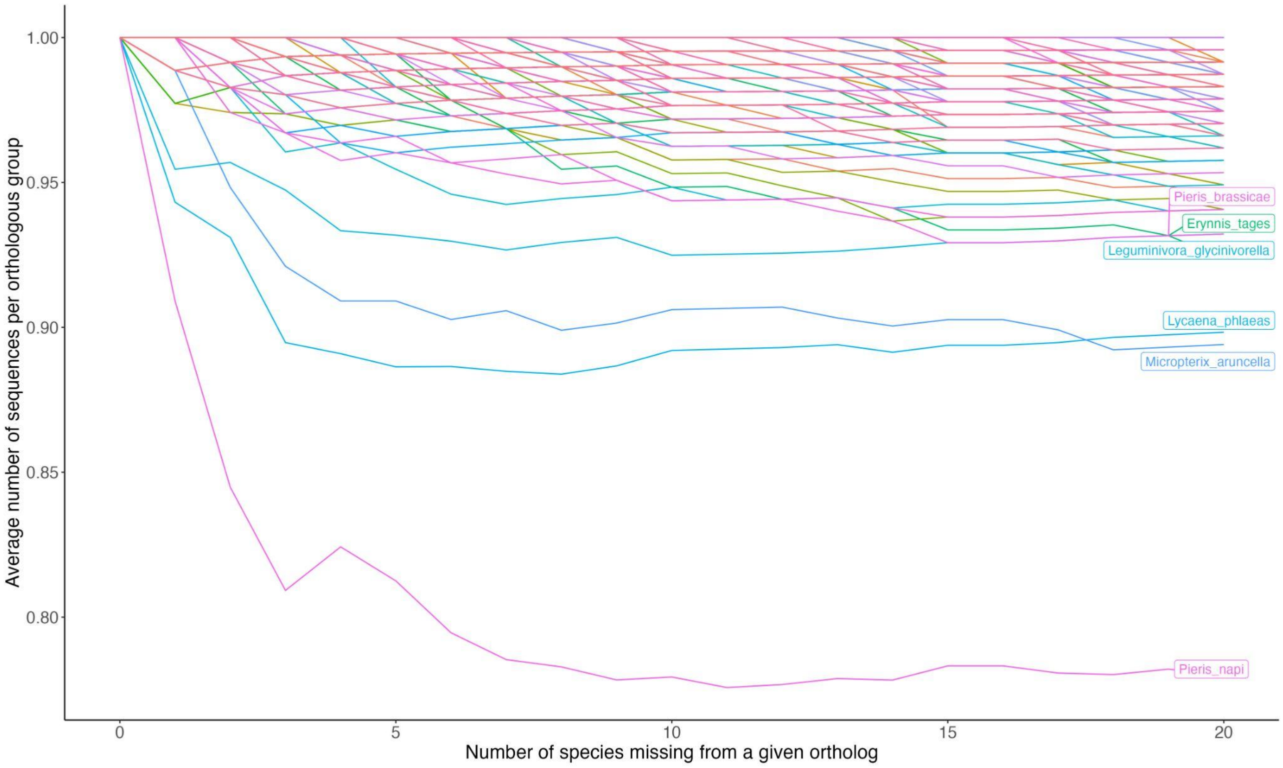


**Fig. S1: Assessing quality of gene annotations using the number of missing conserved single-copy genes**

The average number of sequences per orthologous group. We varied the number of species allowed to be missing from an orthologous group that was otherwise single-copy and conserved in all other species. Species with a low average number of sequences per orthologous group, such as *Pieris napi,* have relatively incomplete gene sets and/or genomes.

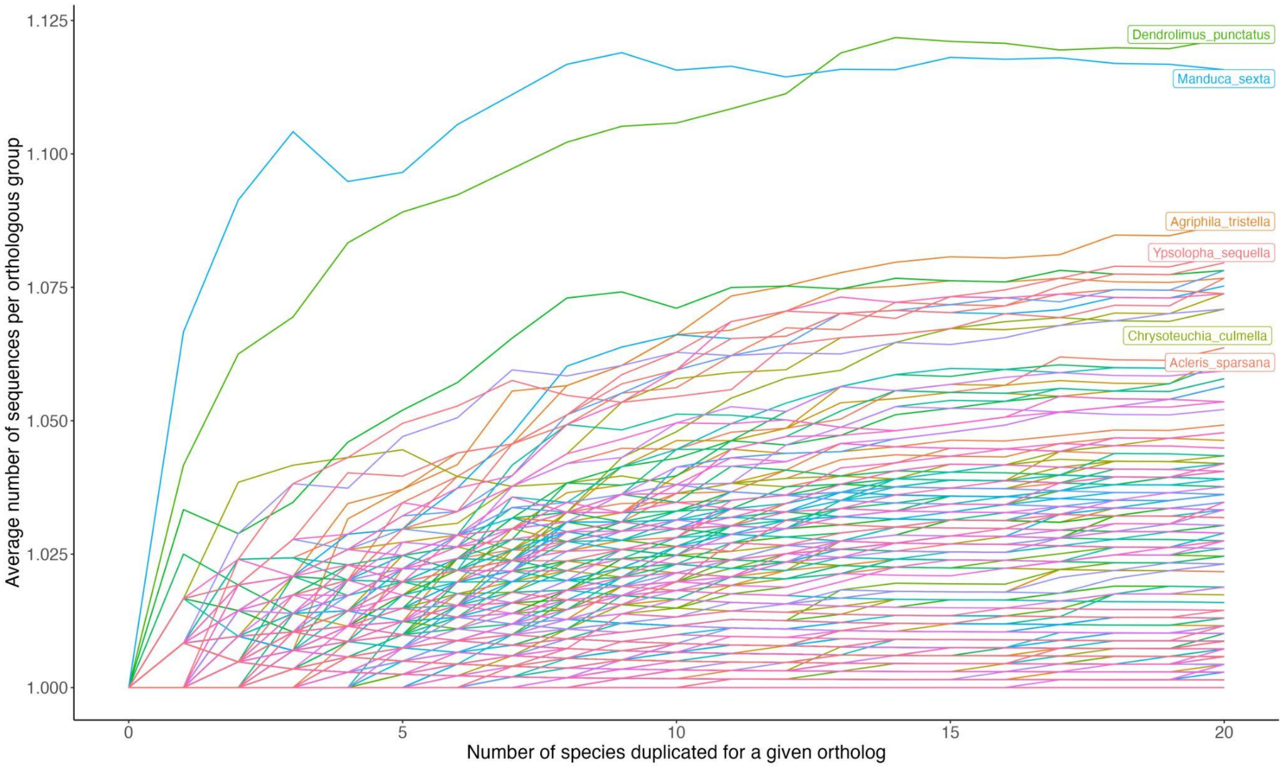


**Fig. S2: Assessing quality of gene annotations using the number of duplicated conserved single-copy genes**

The average number of sequences per orthologous group. We varied the number of species allowed to have two sequences per an orthologous group that was otherwise single-copy and conserved in all other species. Species with a high average number of sequences per orthologous group, such as *Dendrolimus punctutus and Manduca sexta*, have relatively duplicated gene sets and/or genomes.

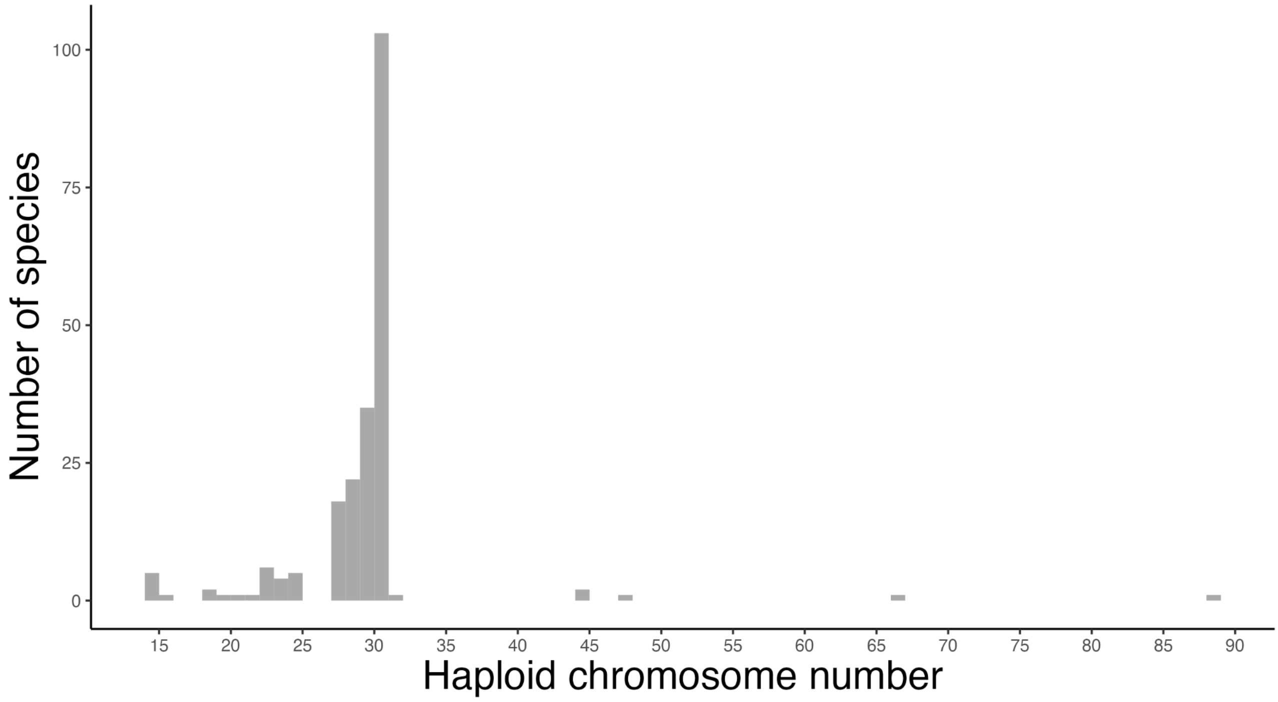


**Fig. S3: Distribution of haploid chromosome number in Lepidoptera from 210 species**
Chromosome numbers obtained from chromosomal genome sequences, including the Z but not W chromosome(s).

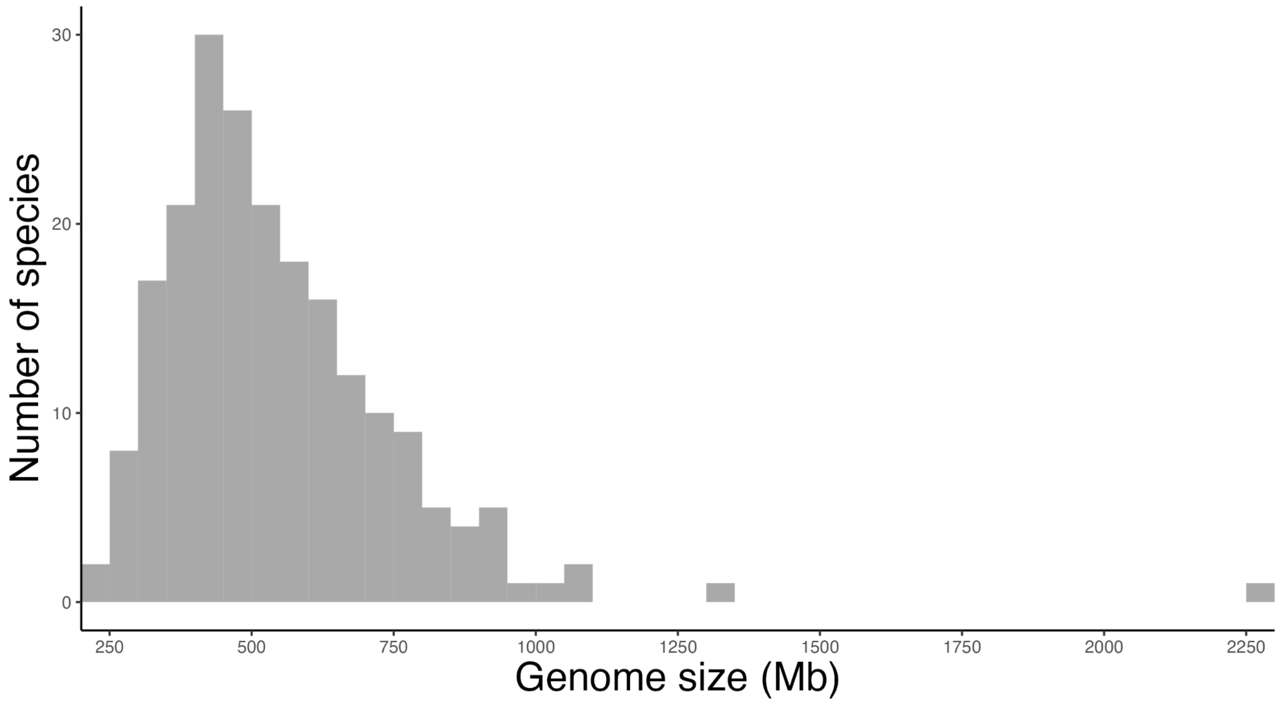


**Fig. S4: Distribution of genome size (Mb) in Lepidoptera from 210 species**
Genome sizes obtained from chromosomal genome sequences.

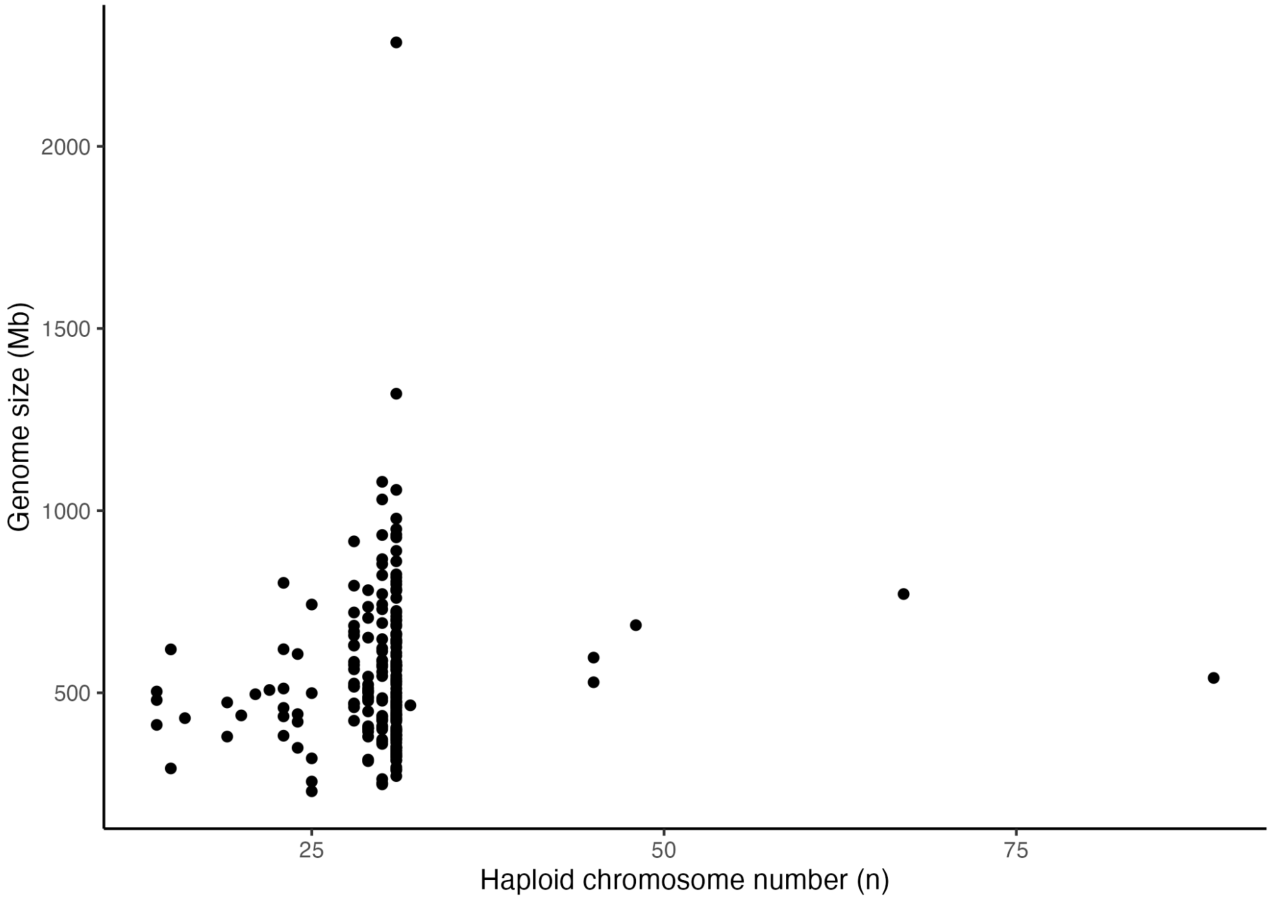


**Fig. S5: Genome size is not correlated with haploid chromosome number**
To account for shared ancestry between species, a phylogenetic linear model was constructed with genome size as the response variable and chromosome number as the predictor variable ($t=0.83$, $p=0.4087$, adjusted $r^2 = 0.00795$). The most appropriate model for the error terms was identified as Ornstein-Uhlenbeck (OU) based on comparison of AIC values. Genome size and chromosome numbers obtained from chromosomal genome sequences.

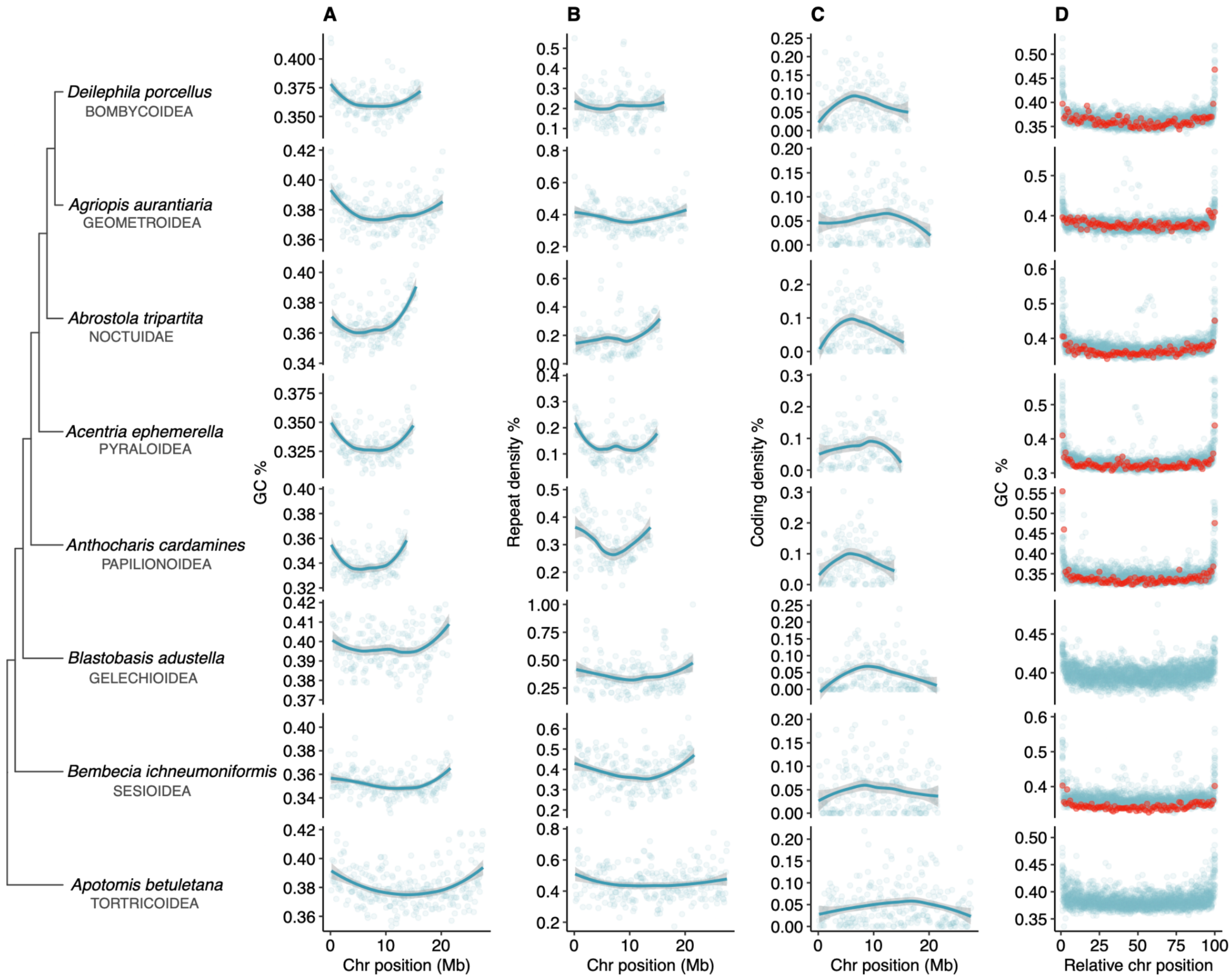


**Fig. S6: Sequence patterns in Lepidopteran chromosomes.**
Sequence patterns for one representative per superfamily in 100 kb non-overlapping windows. (A) GC content in M1 (B) repeat density, considering all repetitive elements, in M1 (C) density of coding sequences in M1. (D) GC content per chromosome in 100 windows per chromosome. Z chromosome values coloured in red. Lines represent LOESS smoothing functions fitted to the data.

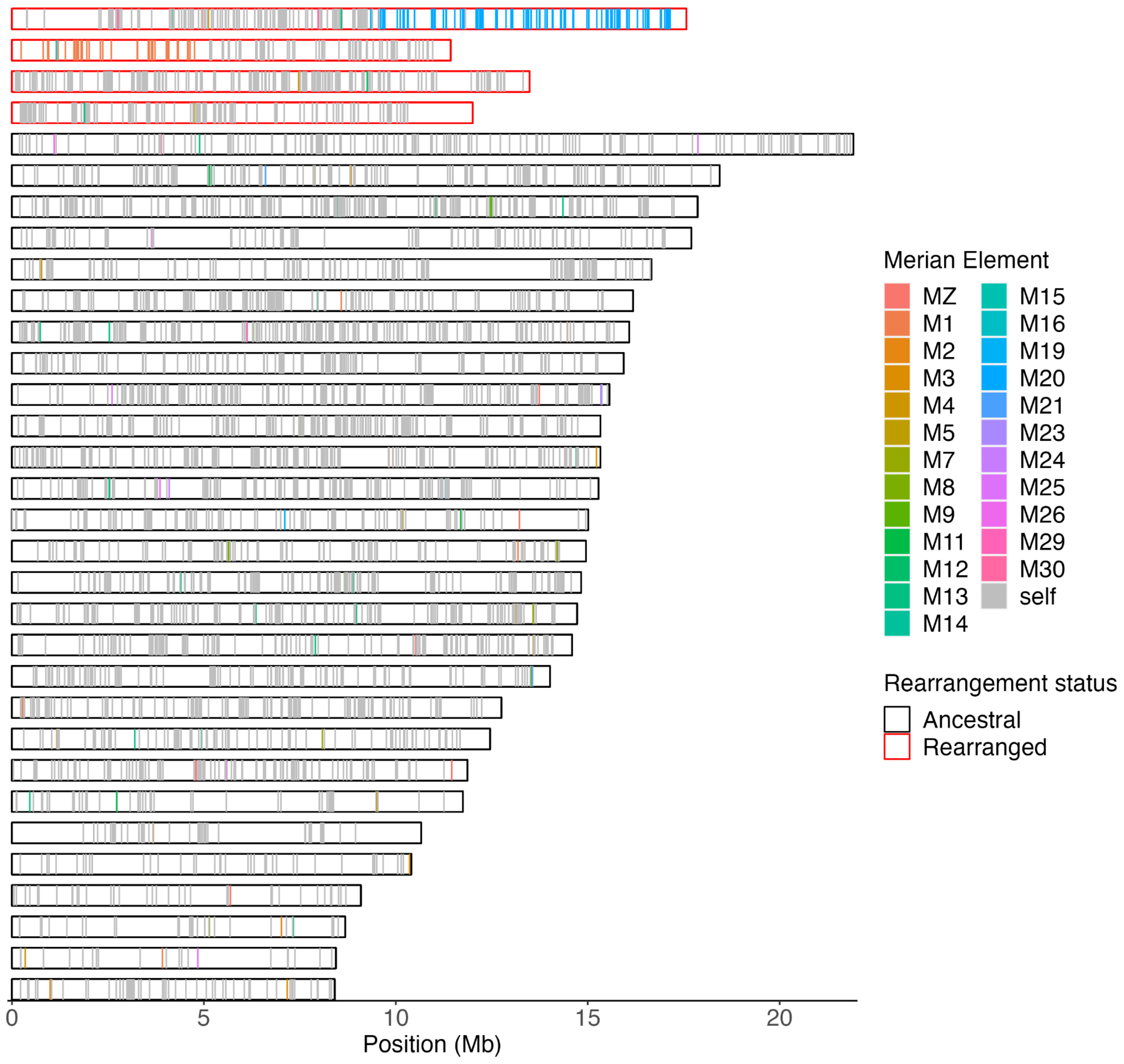


**Fig. S7: Merian elements painted across the chromosomes of *Eupithecia centaureata* demonstrate fission and fusion involving M1 and M6.**
Each chromosome is represented by a rectangle within which the position of each ortholog is grey if it belongs to the most common Merian element for that chromosome or is coloured if it belongs to an alternative Merian element. Chromosomes that have undergone fusions and/or fission events are outlined in red. This reveals a segment of M1 has fused to a segment of M6 (row 2). The remainder of M1 and the remainder of M6 exist as two separate chromosomes (row 3 and 4 respectively), indicating two fission events.

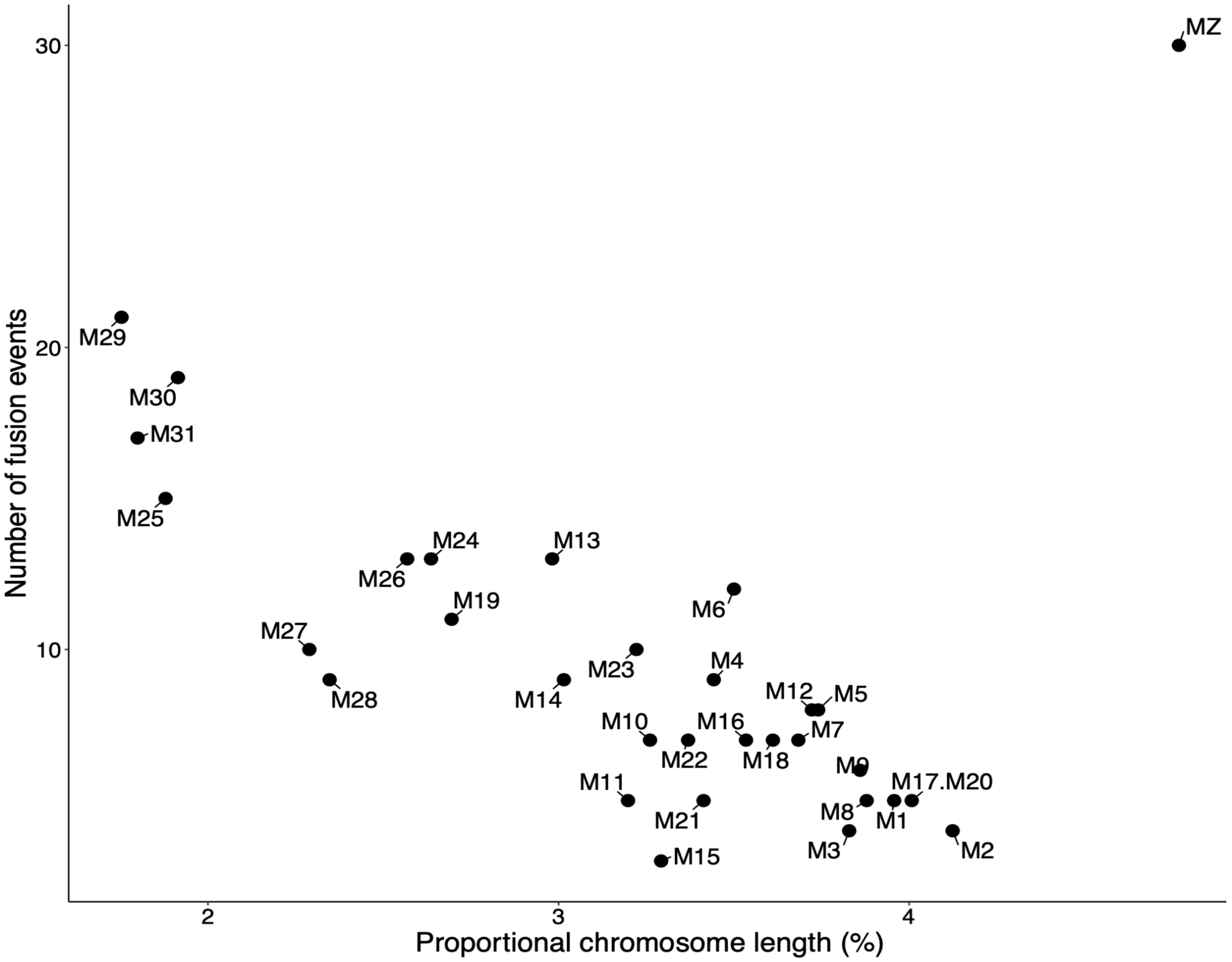


**Fig. S8: Relationship between proportional length and frequency of fusion events**
Number of fusion events that each Merian element is involved in against the average proportional length of the Merian element in the 210 species.

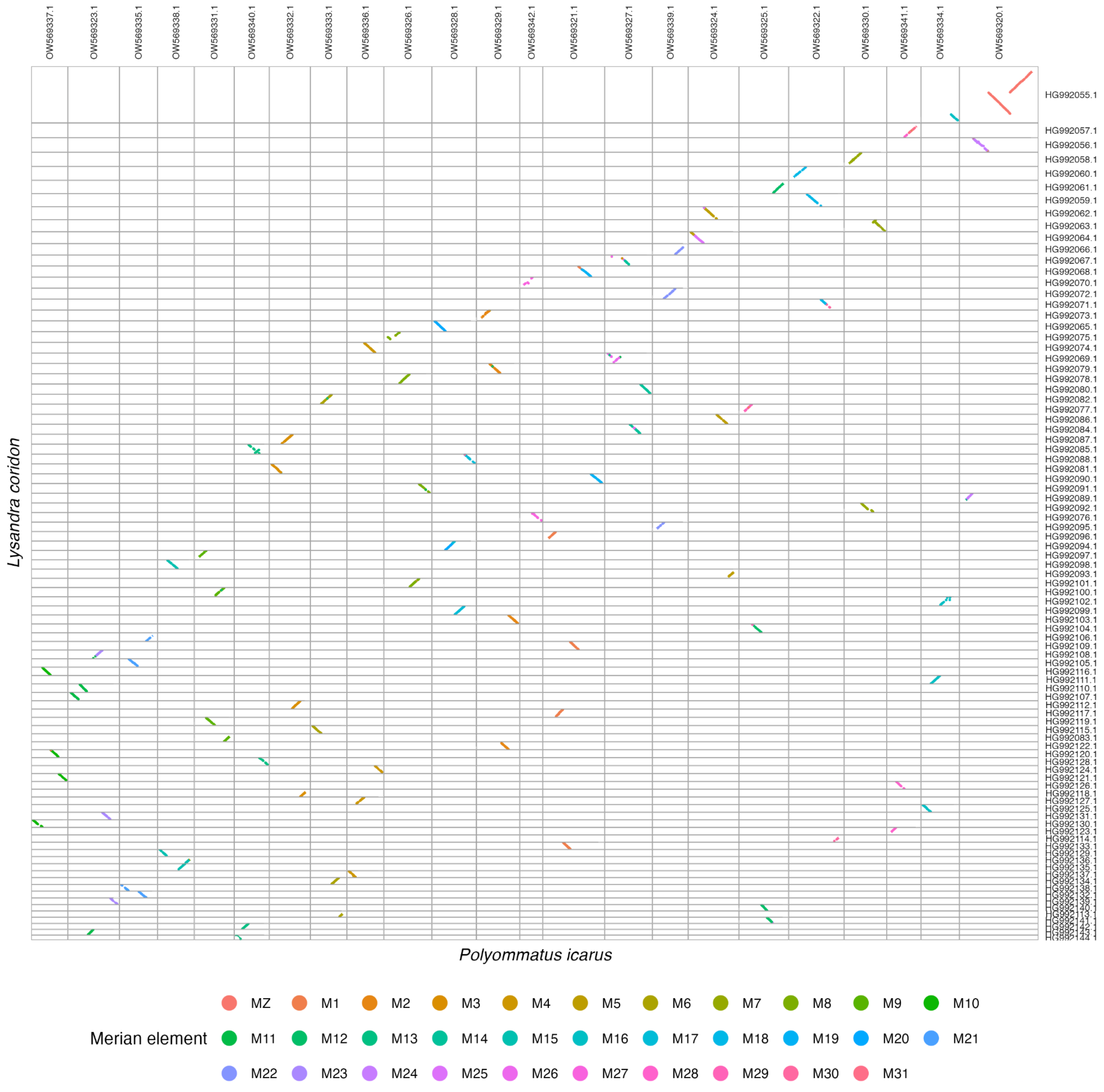


**Fig. S9: Conservation of gene order in *Lysandra coridon* relative to *Polyommatus icarus* despite numerous fission events.**
Oxford plot of relative ortholog positions in the genomes of *Lysandra coridon* and *Polyommatus icarus.* Orthologs are coloured by Merian element.

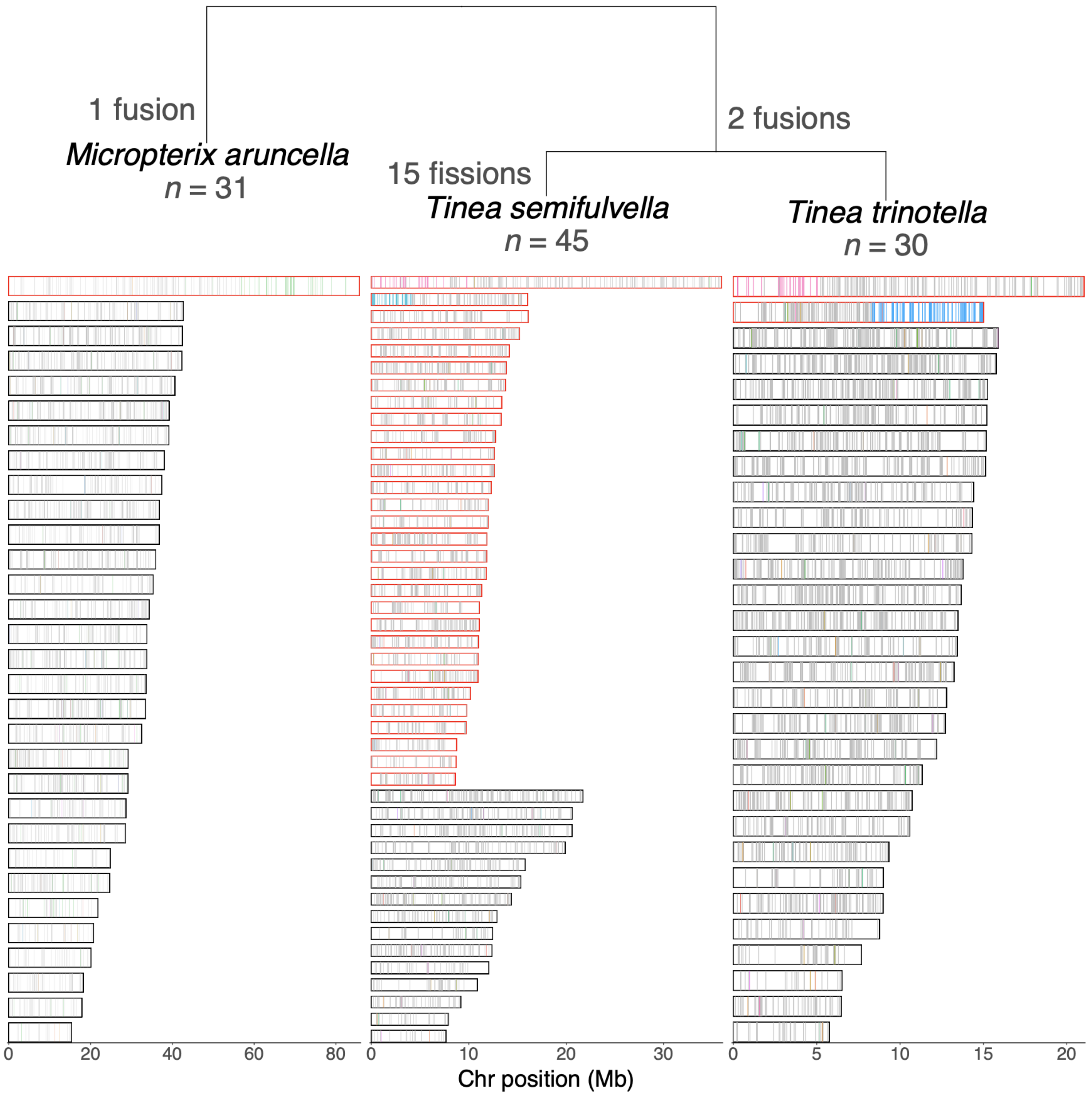


**Fig. S10: Merian elements in two *Tinea* species relative to *Micropterix aruncella***
Relationship between *Tinea semifulvella* and *T. trinotella* and *Micropterix aruncella* annotated with inferred fusion and fission events at each node.

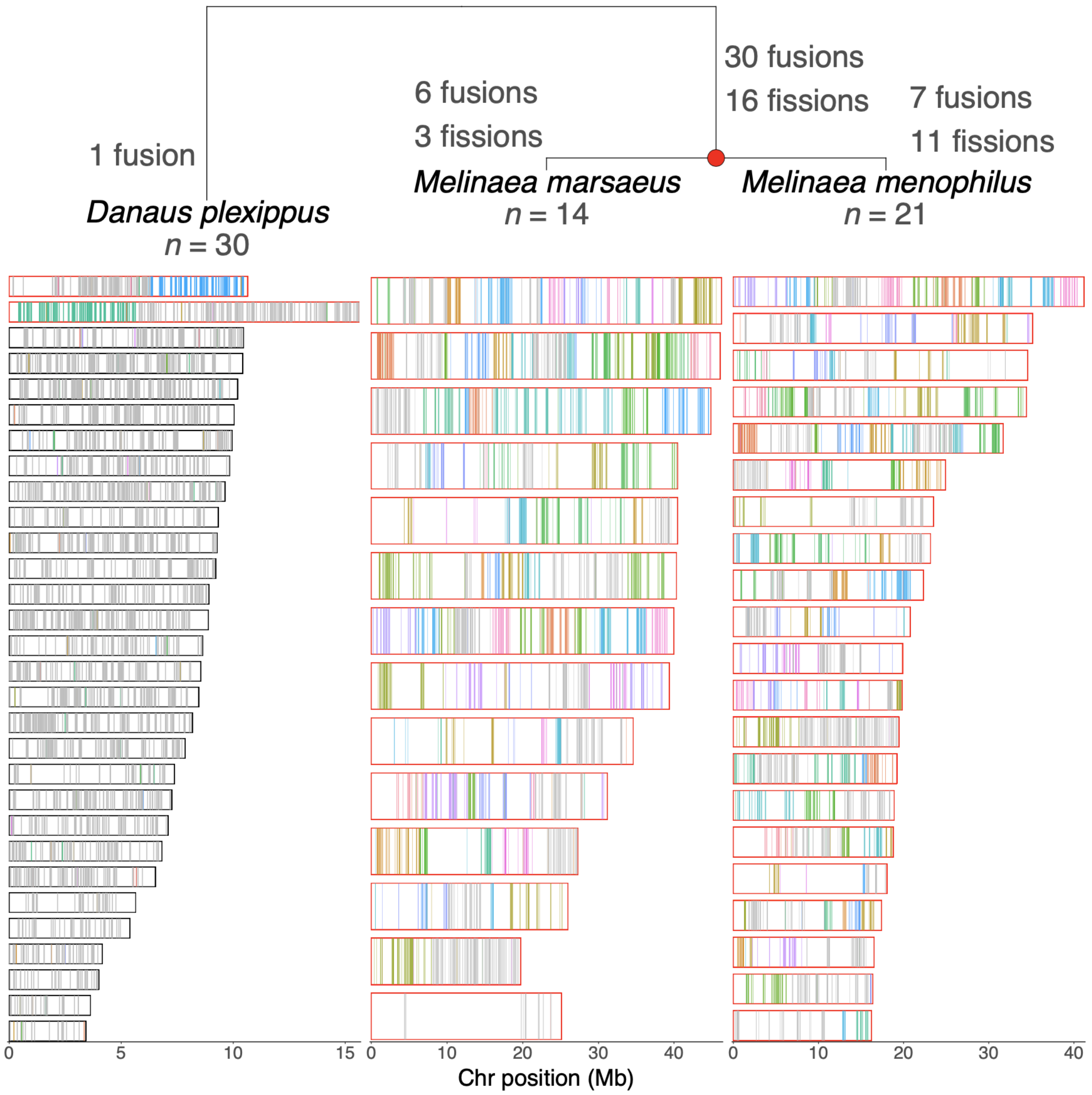


**Fig. S11: Merian elements in two Melinaeae species relative to *Danaus plexippus***
Relationship between *Melinaea marsaeus, M. menophilus* and *Danaus plexippus* annotated with inferred fusion and fission events at each node.

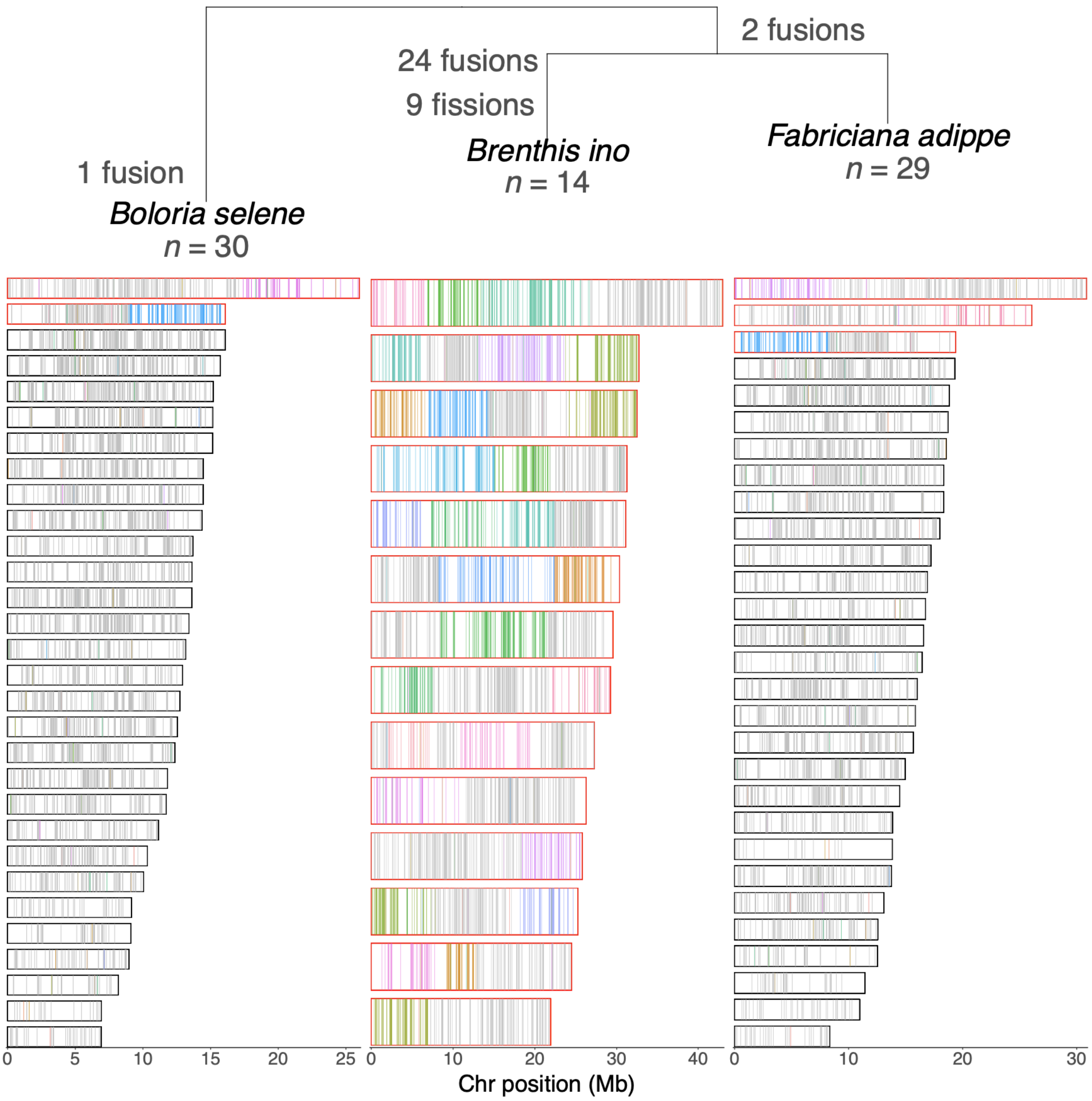


**Fig. S12: Merian elements in *Brenthis ino* relative to *Fabriciana adippe* and *Boloria selene.*** Relationships between *Brenthis ino, Fabriciana adippe*a and *Boloria selene* annotated with inferred fusion and fission events at each node.

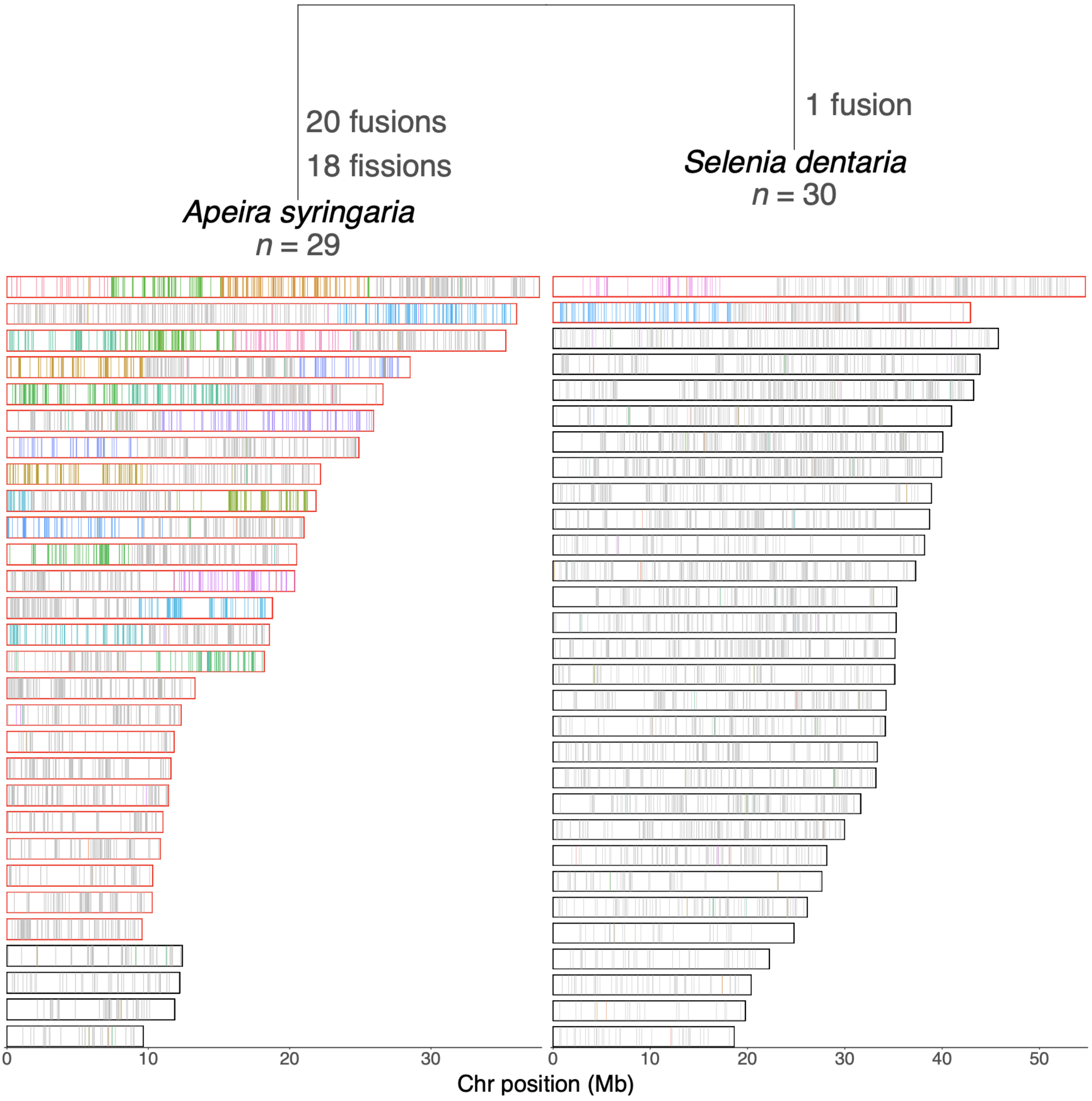


**Fig. S13: Merian elements in *Apeira syringaria* relative to *Selenia dentaria***
Relationship between *Apeira syringaria* and *Selenia dentaria* annotated with inferred fusion and fission events at each node.

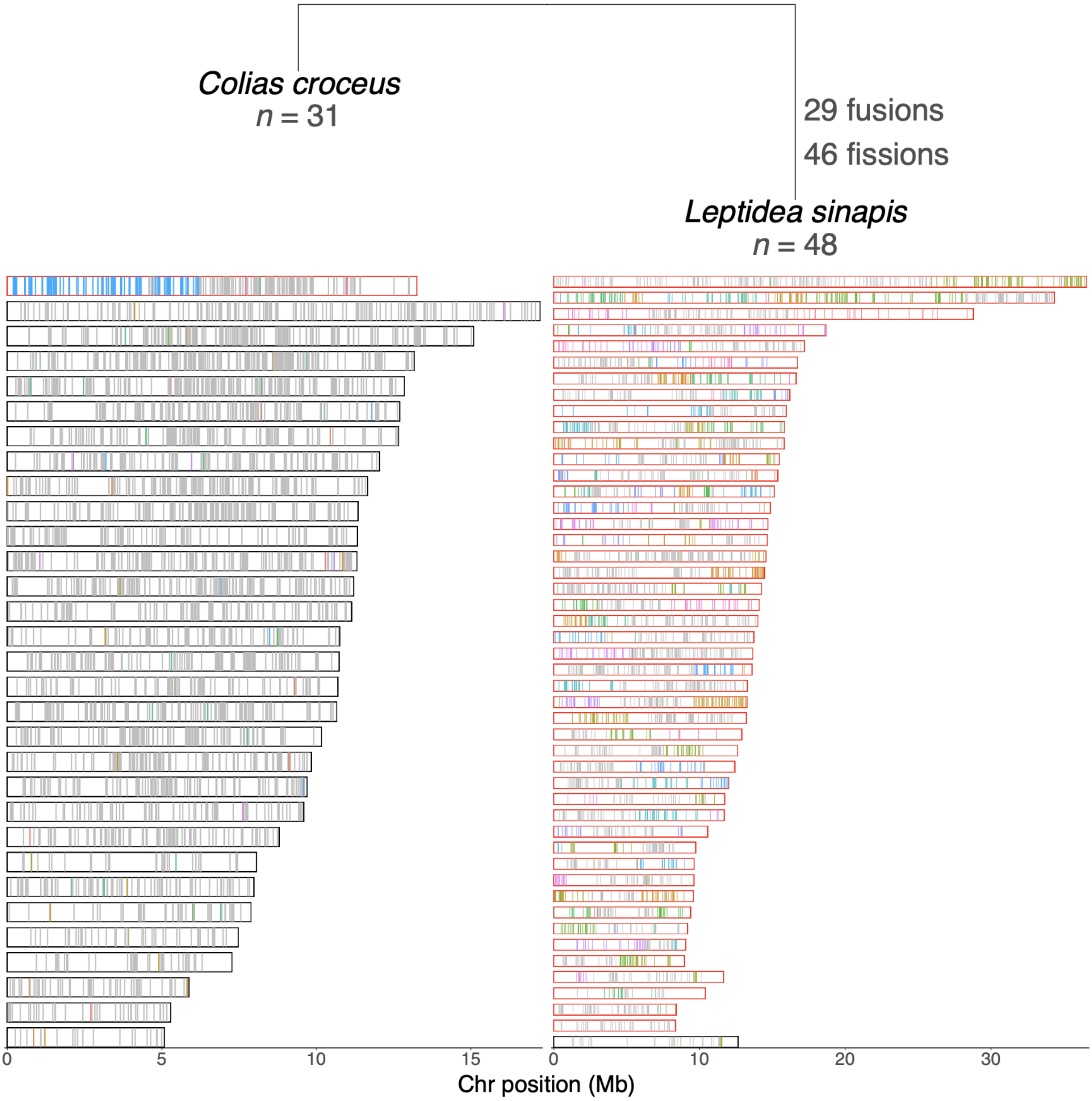


**Fig. S14: Merian elements in *Leptidea sinapis* relative to *Anthocharis cardamines***
Relationships between *Colias croceus* and *Leptidea sinapis* annotated with inferred fusion and fission events at each node.

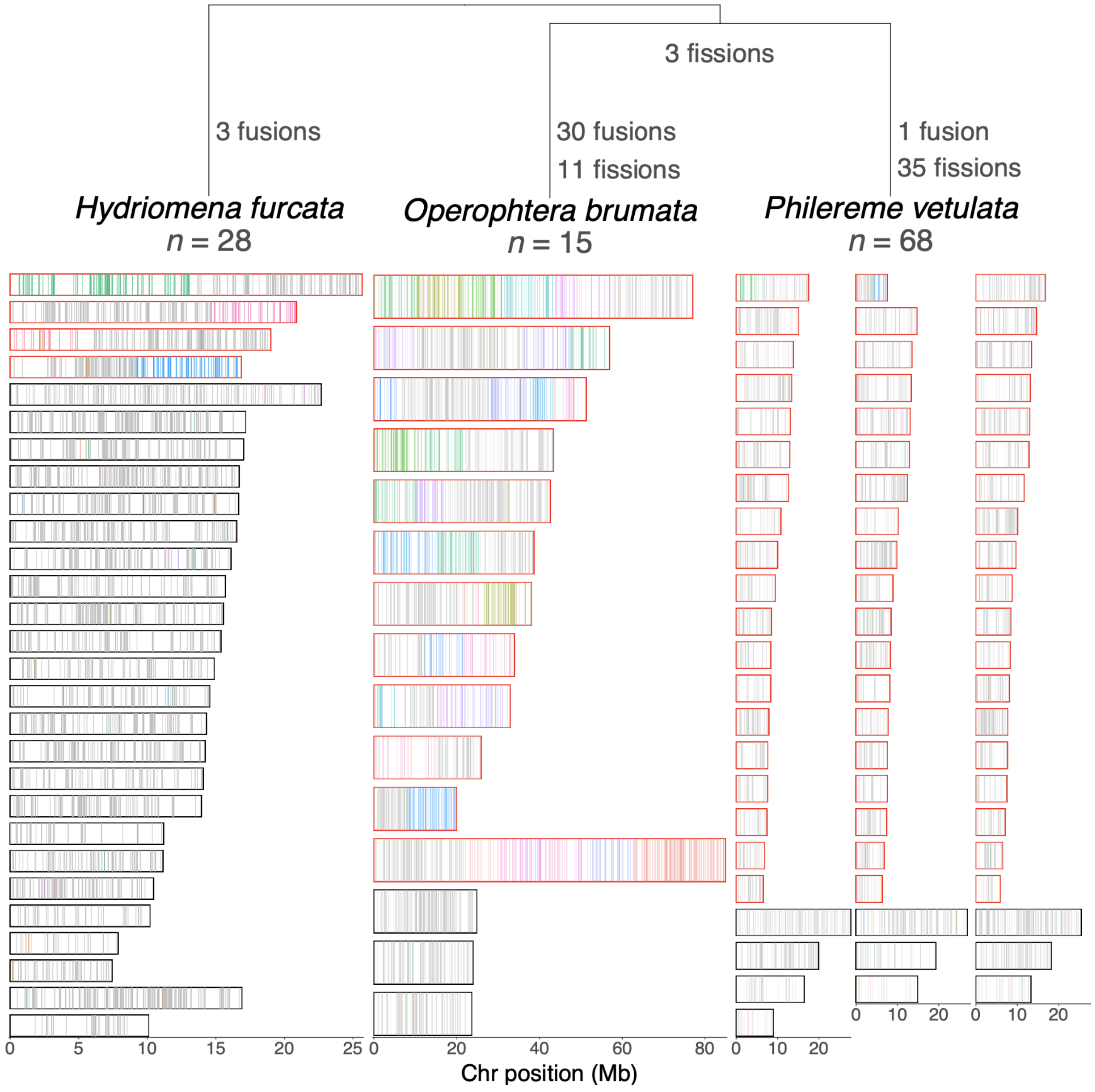


**Fig. S15: Merian elements in *Operophtera brumata* and *Philereme vetulata* relative to *Hydriomena furcata***
Relationships between *Hydriomena furcata*, *Operophtera brumata* and *Philereme vetulata* annotated with inferred fusion and fission events at each node.

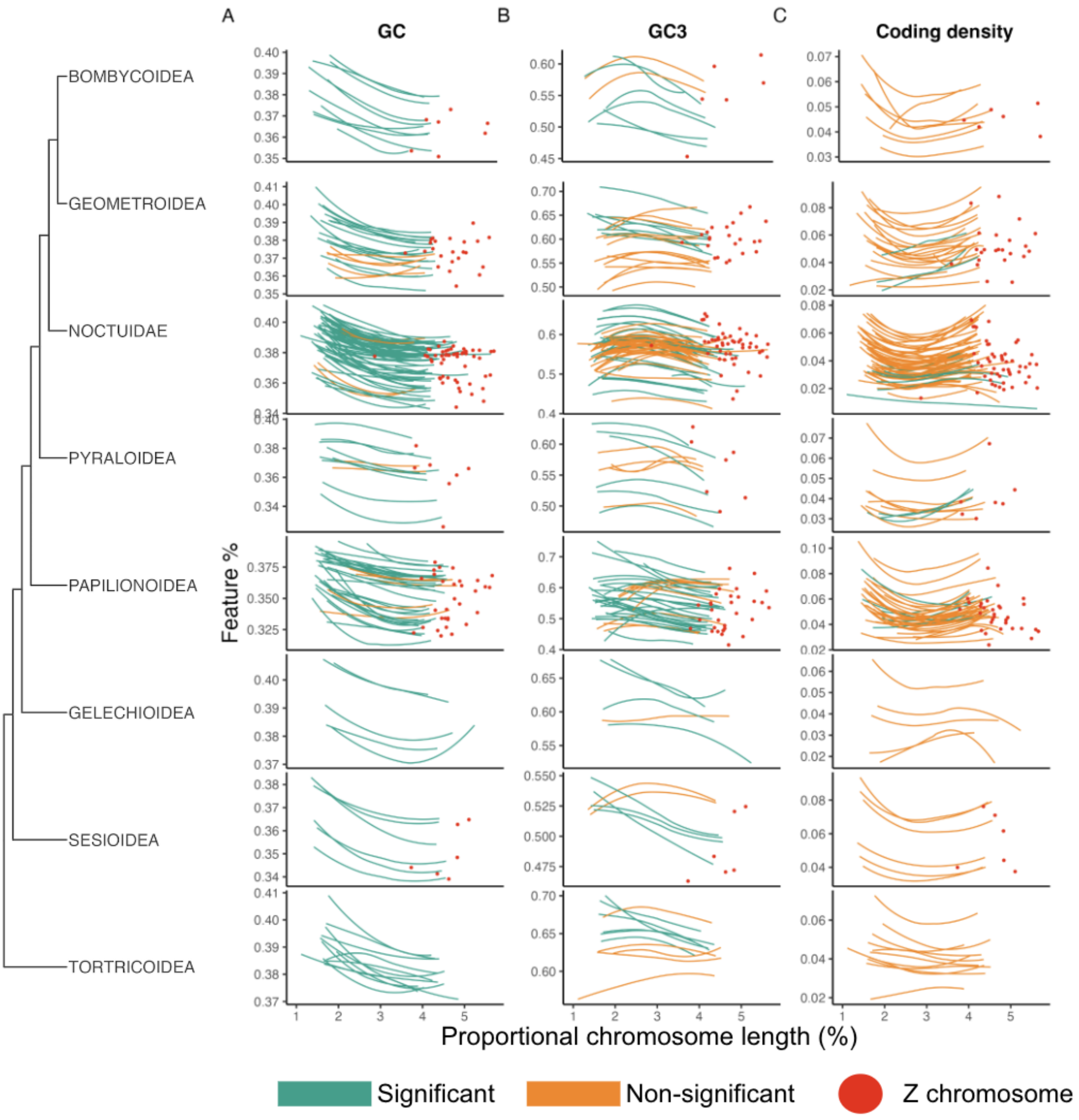


**Fig. S16: Correlation of GC, GC3 and coding density with chromosome length.**
Proportional chromosome length against (A) GC proportion, (B) GC percentage at the third codon position (GC3), (C) density of coding sequence. Proportional chromosome length is chromosome length (Mb) divided by genome size per species. A locally weighted smoothing line ("loess") is drawn between the autosomes of each species. The line is coloured green if the correlation was significant (Spearman's rank, $p < 0.05$), or orange if it was non-significant. Only species with at least 10 autosomes are included and all autosomes which are inferred to have undergone fusion or fission events were removed. Only superfamilies represented by at least 5 superfamilies are shown. The Z chromosomes were not included in the correlation analysis and are drawn in red.

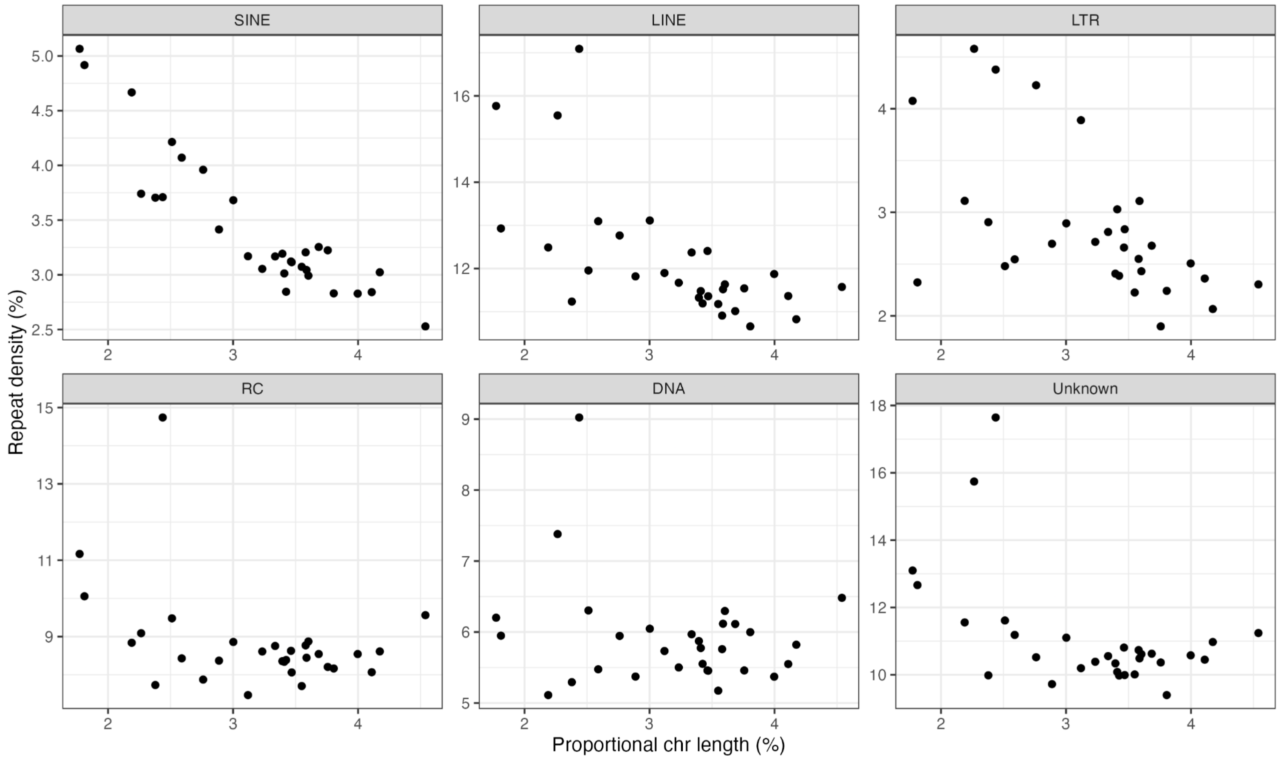


**Fig. S17: Relationship between repetitive element density and proportional chromosome length for each major class of transposable elements**

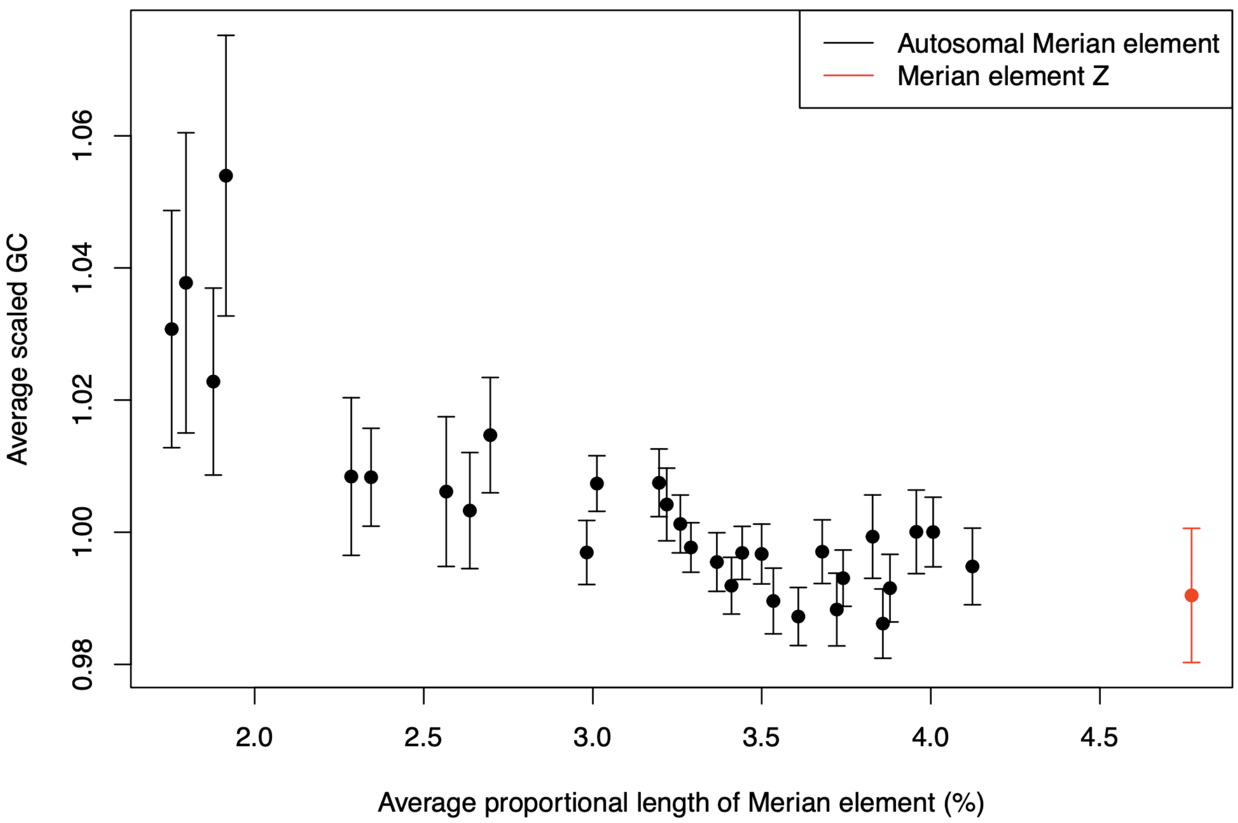


**Fig S18: Negative relationship between GC and proportional chromosome length of each Merian element**

Mean proportional chromosome length (length divided by genome size) per Merian element against the mean GC value (%) of the Merian element scaled by mean GC content of the genome. Merian element Z is indicated in red.

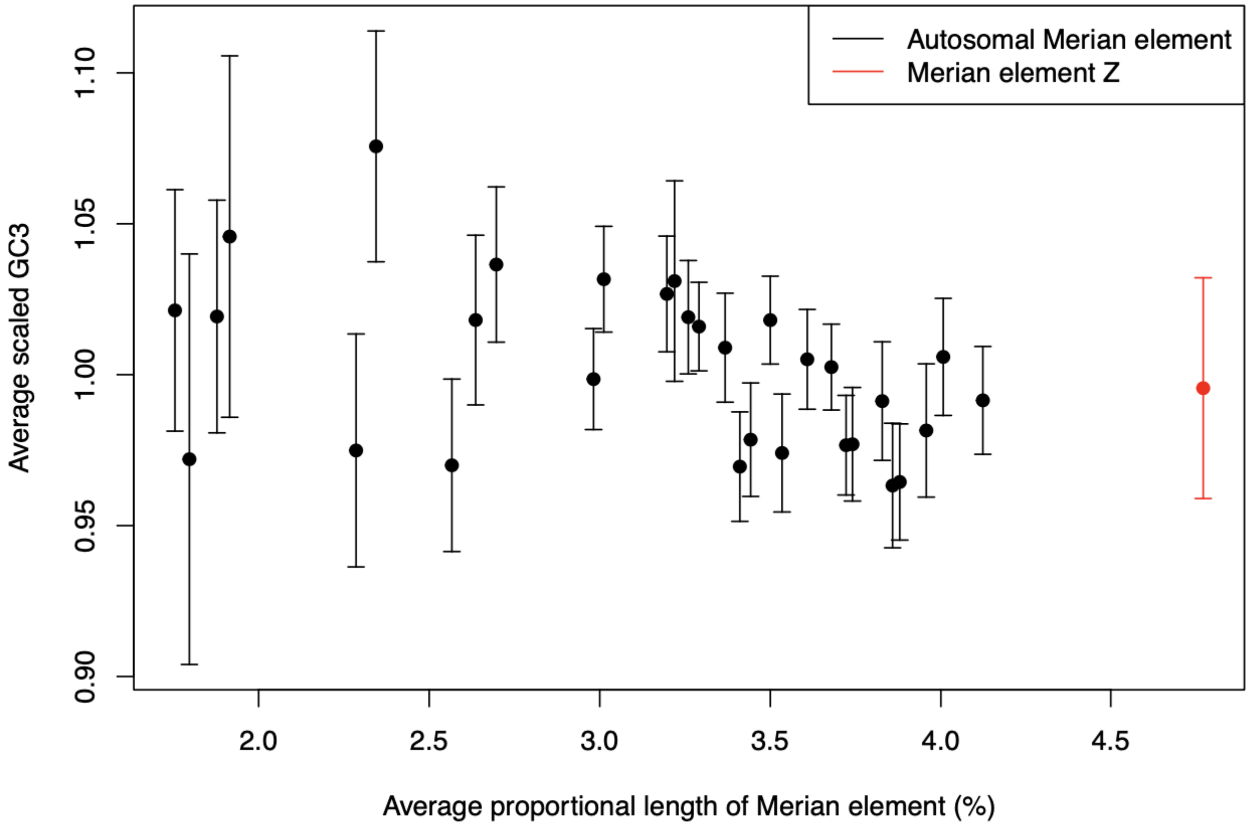


**Fig S19: Negative relationship between GC3 and proportional chromosome length of each Merian element**

Mean proportional chromosome length (length divided by genome size) per Merian element against the mean GC3 value (%) of the Merian element scaled by mean GC3 content of the genome. Merian element Z is indicated in red.

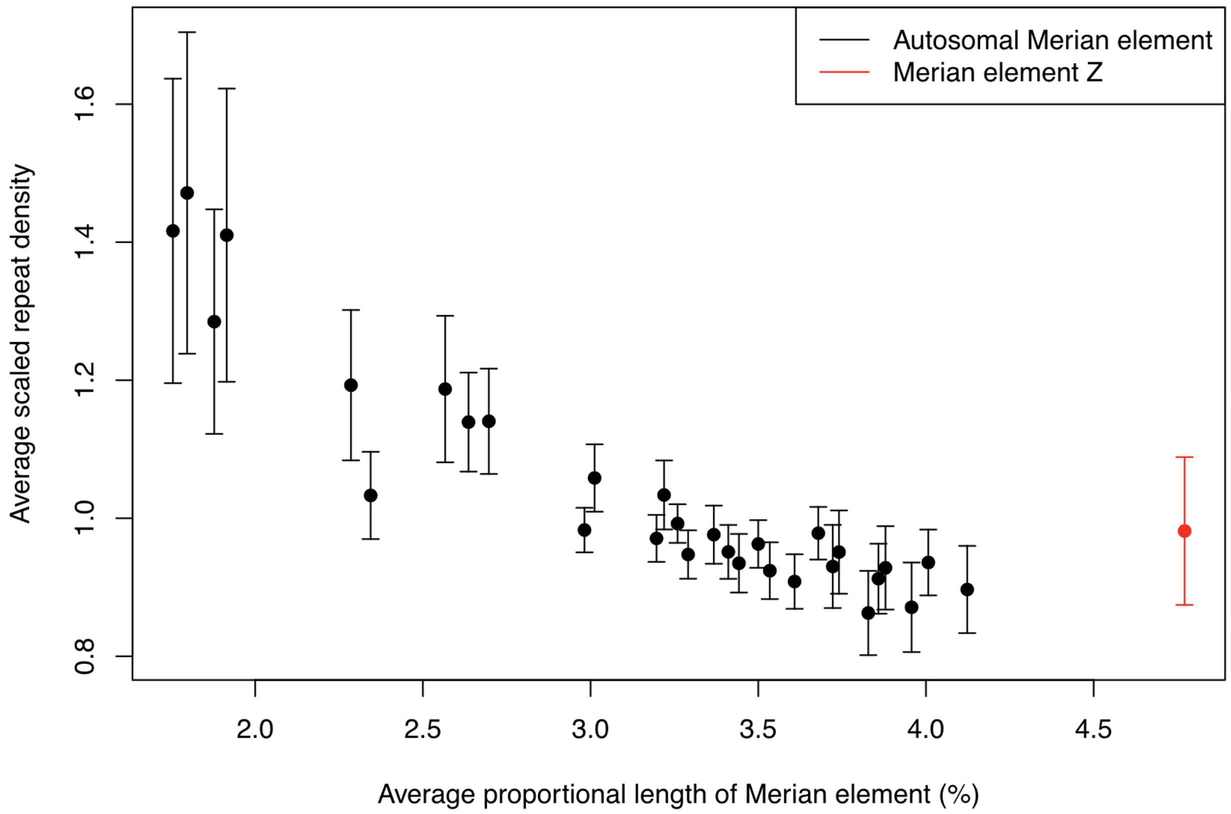


**Fig S20: Negative relationship between repeat density and proportional chromosome length of each Merian element**

Mean proportional chromosome length (length divided by genome size) per Merian element against the mean repeat density of the Merian element scaled by mean repeat density of the genome. Merian element Z is indicated in red.

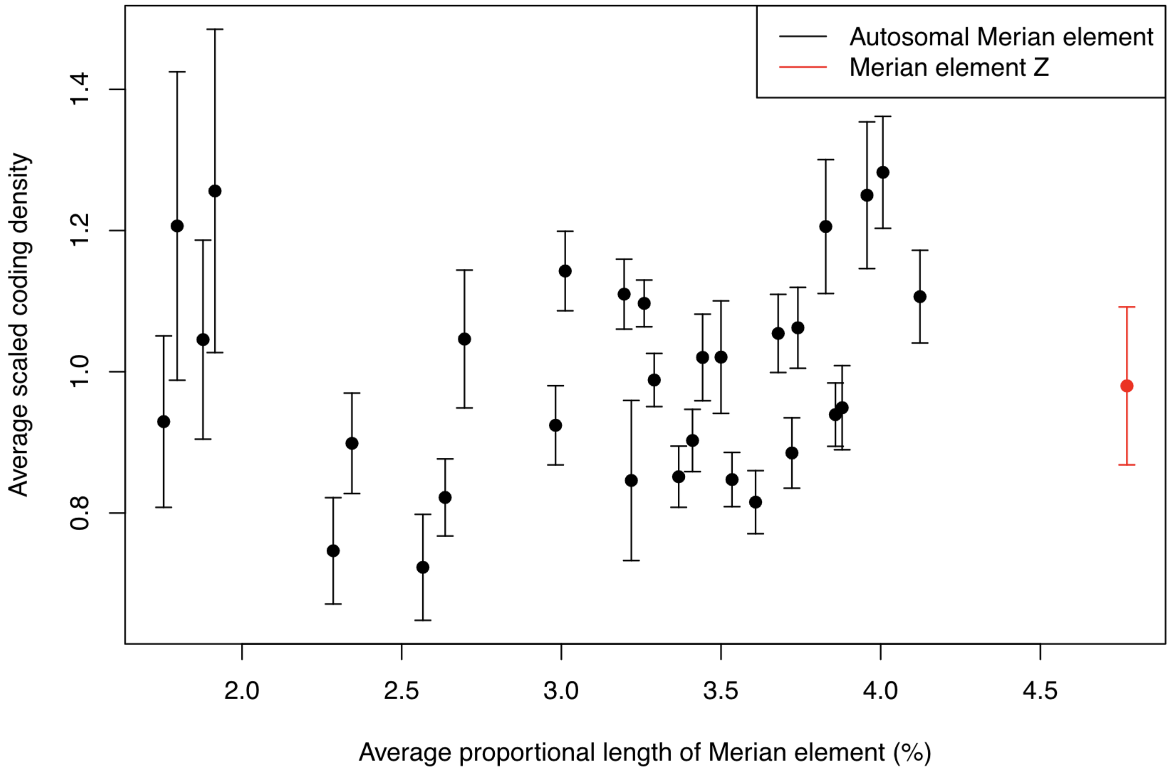


**Fig S21: Variation in the coding density against proportional chromosome length of each Merian element**

Mean proportional chromosome length (length divided by genome size) per Merian element against the mean coding density of the Merian element scaled by mean coding density of the genome. Merian element Z is indicated in red.

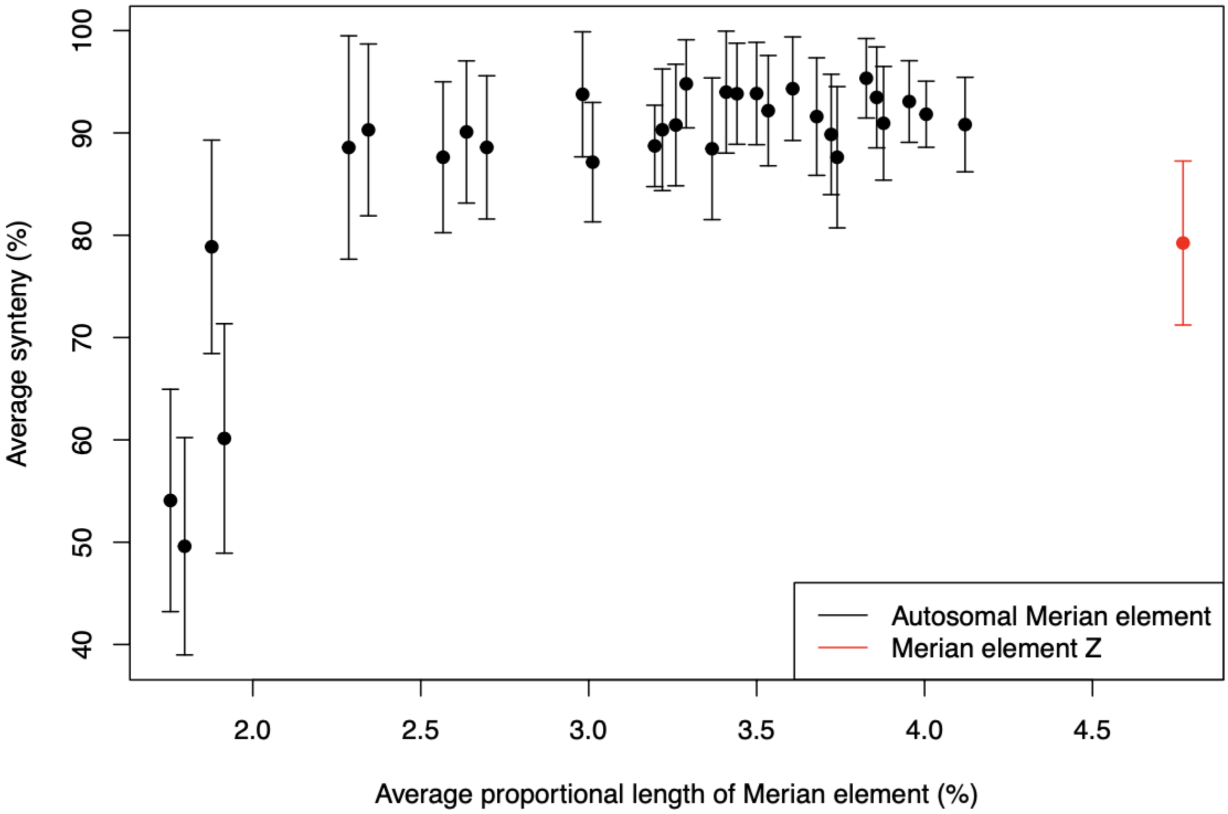


**Fig S22: Positive relationship between the level of synteny and proportional chromosome length of each Merian element**

Mean proportional chromosome length (length divided by genome size) per Merian element against the mean level of synteny of the Merian element. Merian element Z is indicated in red. Only chromosomes which have not undergone fusions or fission events are included.

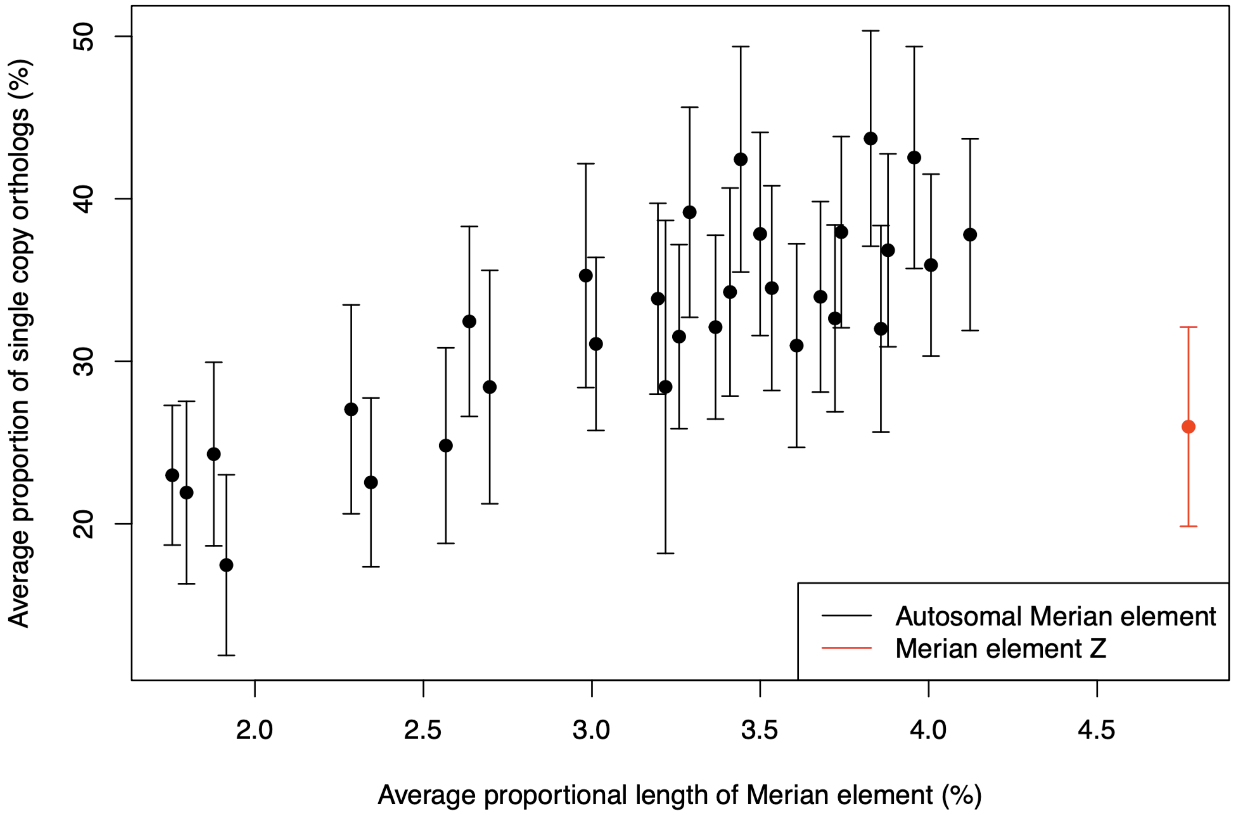


**Fig S23: Positive relationship between the proportion of single copy orthologs and proportional chromosome length of each Merian element**

Mean proportional chromosome length (length divided by genome size) per Merian element against the mean proportion of genes on the chromosome which are single copy and present in the majority of species. Merian element Z is indicated in red. Only chromosomes which have not undergone fusions or fission events are included.

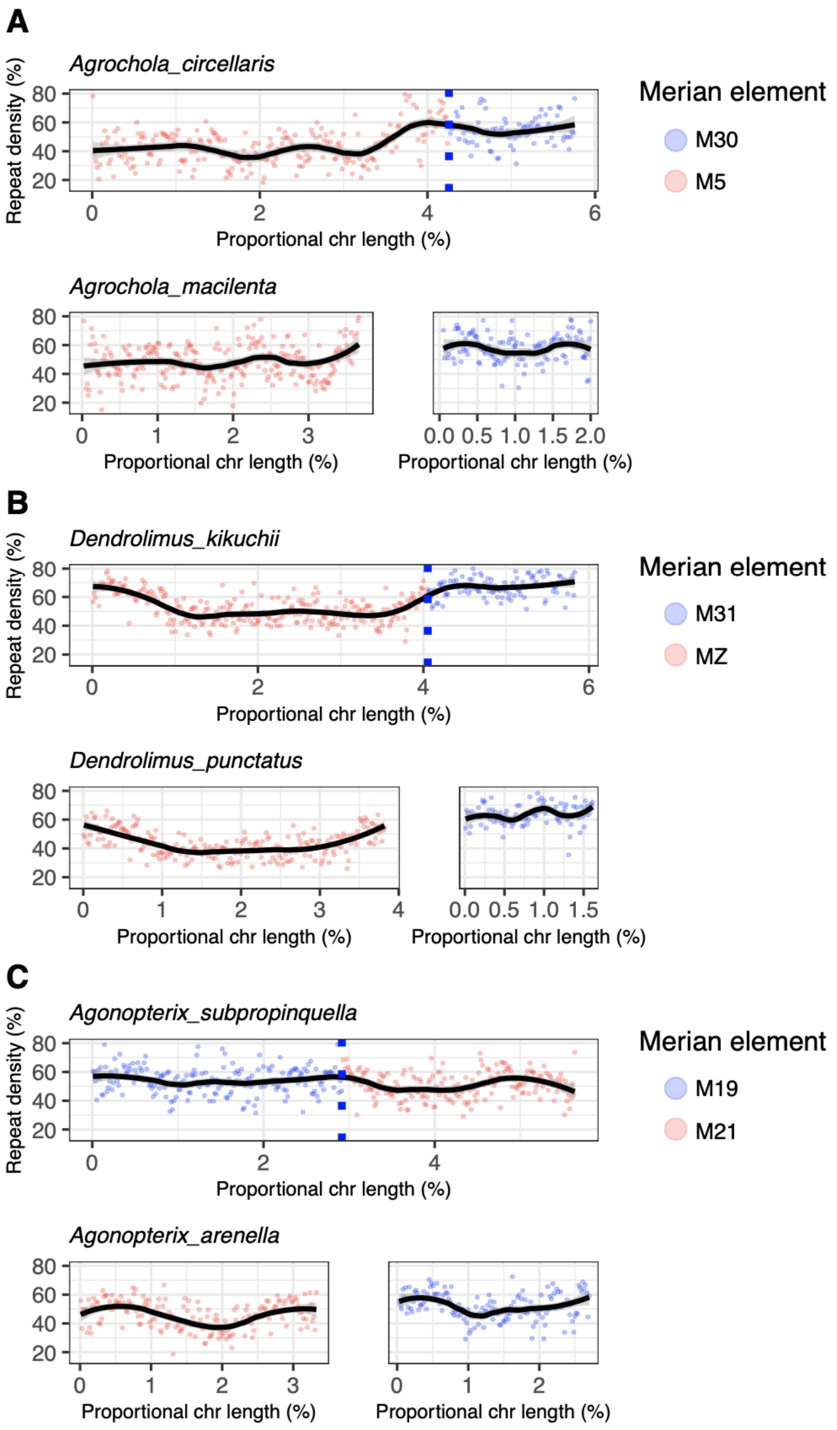


**Fig S24: Repeat density across fused chromosomes and their unfused homologs in pairs of sister species.**

Repeat density in fused chromosomes in one species, compared to the repeat density of the unfused orthologous chromosomes in a sister species. As each fusion is present in one species, and absent in the other sampled species from the same genus, they are likely very recent fusions. (A) *Agrochola circellaris*, compared to *A. macilenta*, (B) *Dendrolimus kikuchii* compared to *D. kikuchii* and (C) *Agonopterix subpropinquella* and *A. arenella.* Repeat density is plotted along each chromosome in 100 kb windows, where the chromosome position is scaled to proportional length by dividing by genome size. Lines represent LOESS smoothing functions fitted to the data. Points are coloured by which Merian element they represent. Blue dashed lines indicate the fusion point along fused chromosomes.

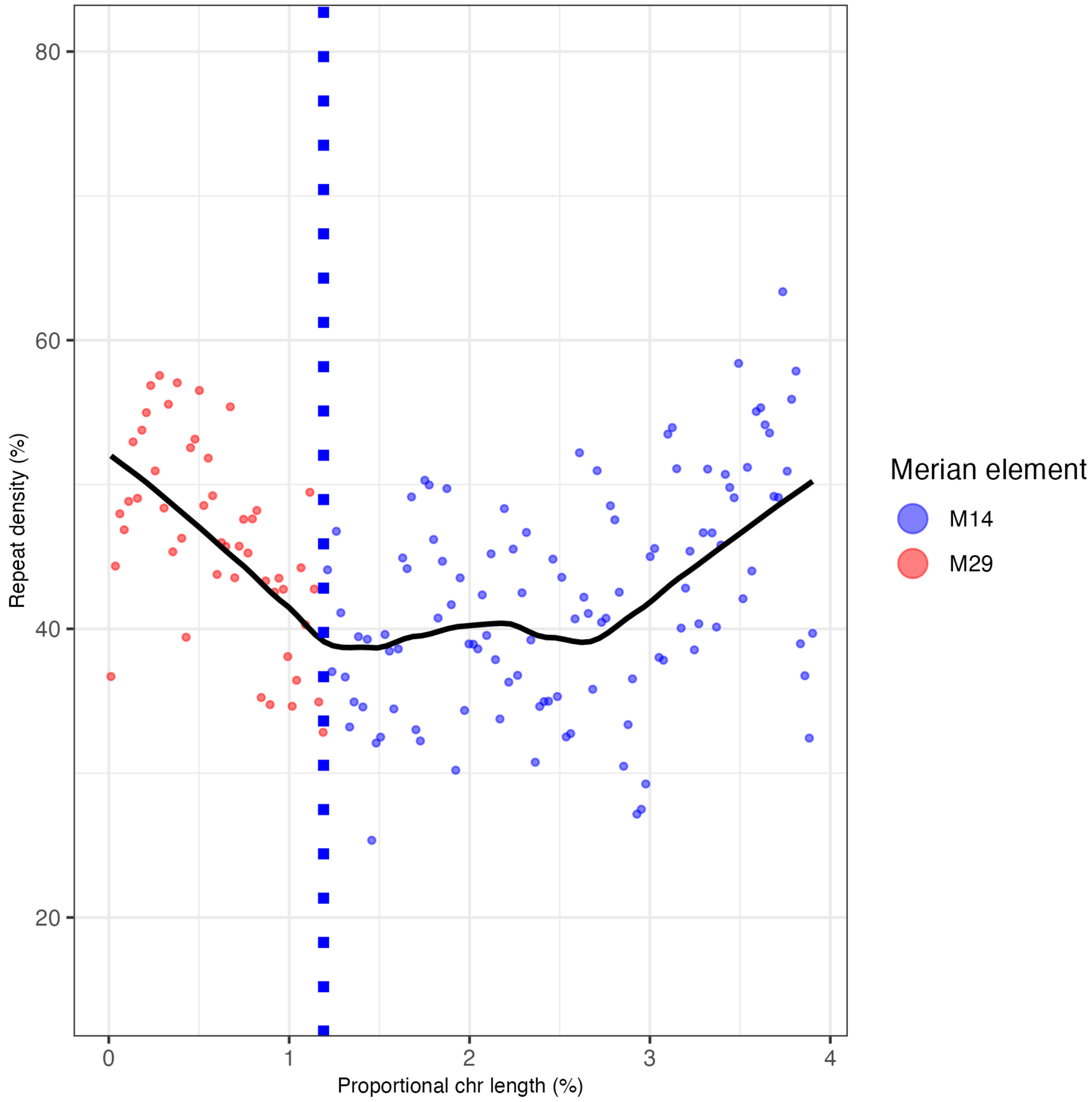


**Fig S25: Repeat density across the fused chromosome in *Aphantopus hyperantus***

Repeat density across the fused chromosome (M29+M14) in *Aphantopus hyperantus.* The distance between *A. hyperantus* and the closest relative with an unfused M29 and M14 (*Danaus plexippus*) is too large for the unfused chromosomes to be a suitable proxy for the ancestral chromosomes and so are not shown. Repeat density is plotted along the chromosome in 100 kb windows, where the chromosome position is scaled to proportional length by dividing by genome size. Lines represent LOESS smoothing functions fitted to the data. Points are coloured by which Merian element they represent. Blue dashed lines indicate the fusion point along fused chromosomes.

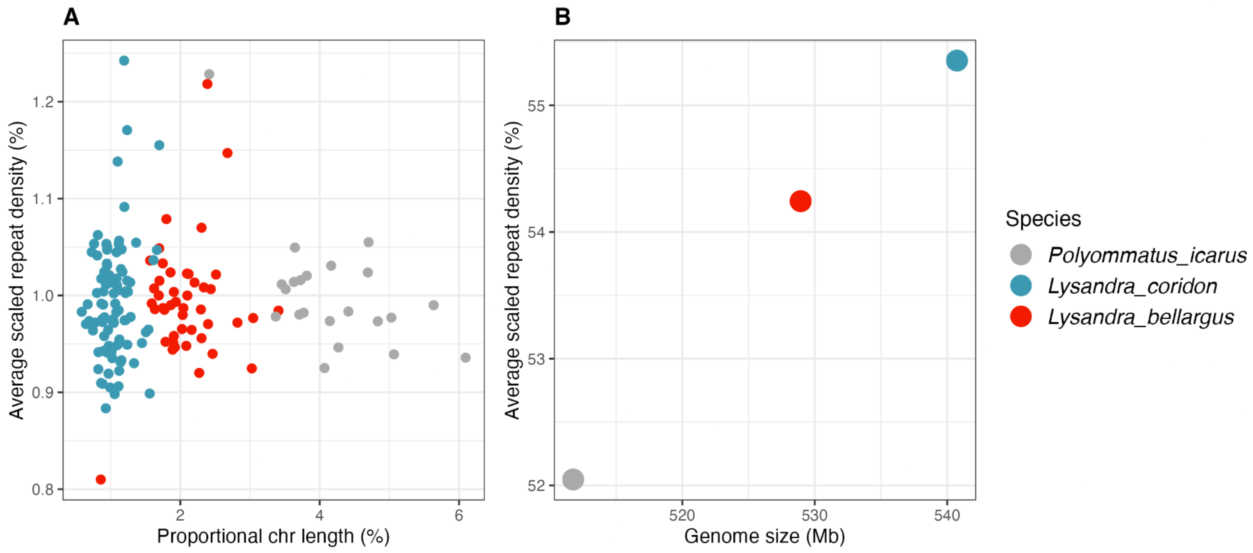


**Fig S26: Relationship between repeat density, genome size and chromosome size in *Lysandra***

Mean scaled repeat density (%) against (A) proportional chromosome length (%) and (B) genome size (Mb).

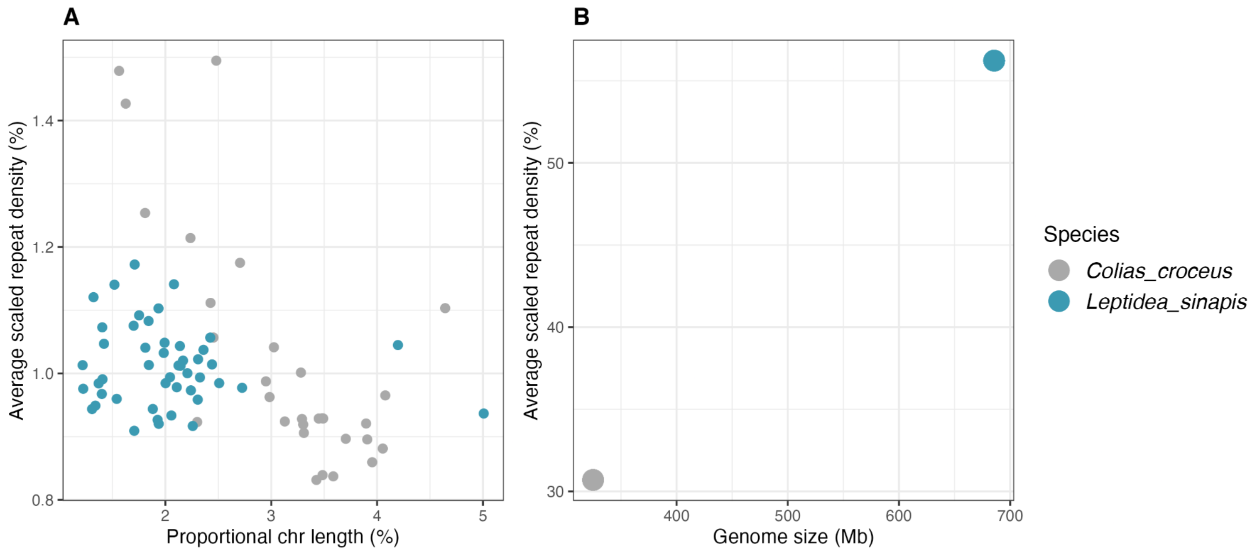


**Fig S27: Relationship between repeat density, genome size and chromosome size in *Leptidea***

Mean scaled repeat density (%) against (A) proportional chromosome length (%) and (B) genome size (Mb).

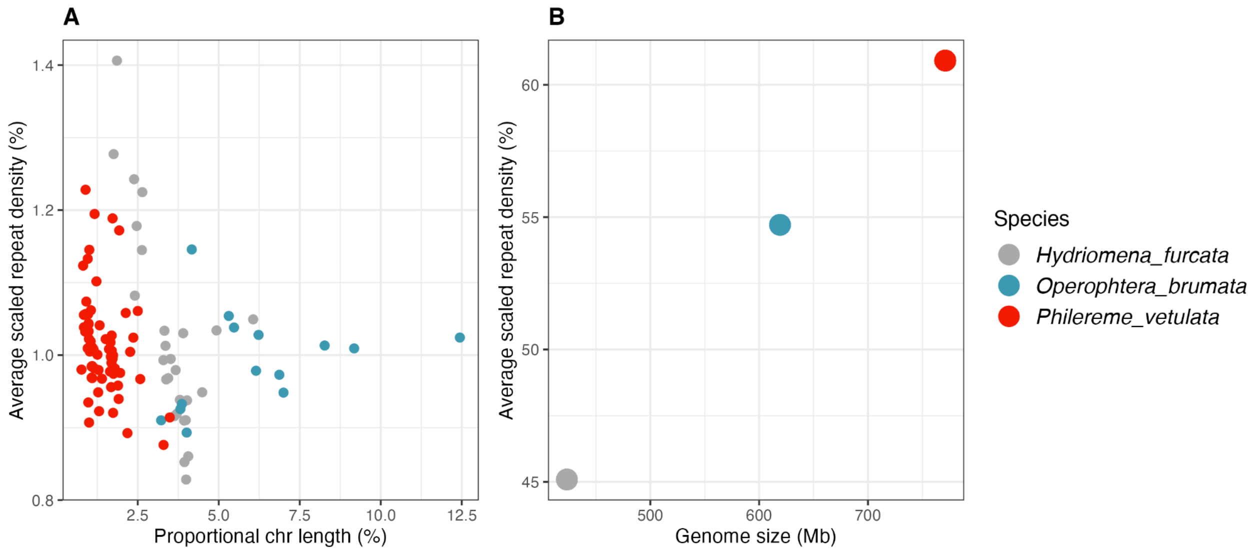


**Fig S28: Relationship between repeat density, genome size and chromosome size in *Philereme vetulara* and *Operophtera brumata***

Mean scaled repeat density (%) against (A) proportional chromosome length (%) and (B) genome size (Mb).

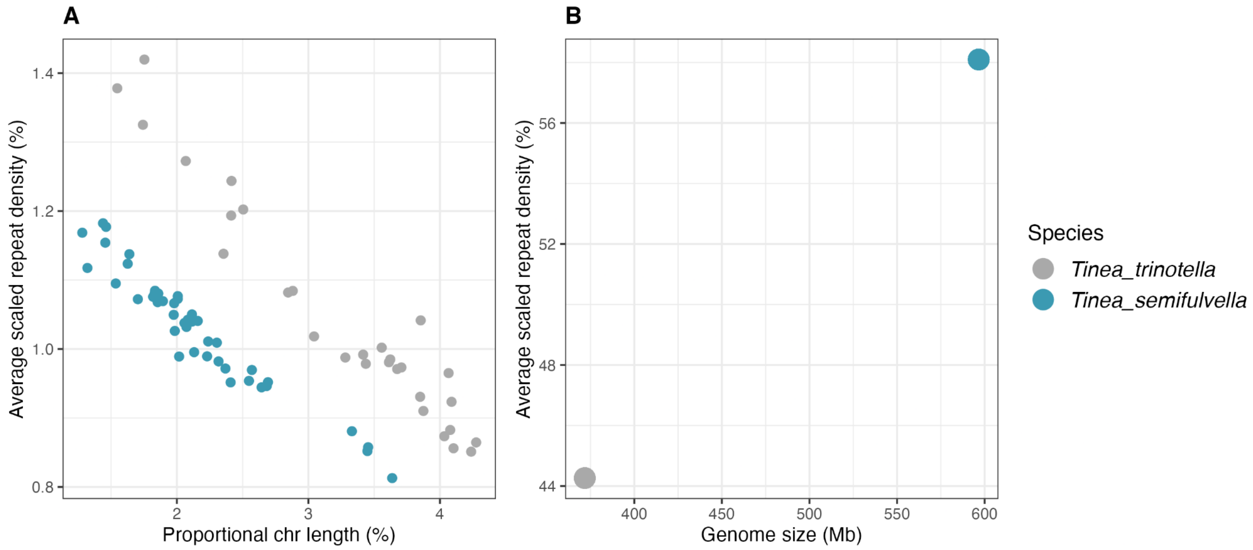


**Fig S29: Relationship between repeat density, genome size and chromosome size in *Tinea***
Mean scaled repeat density (%) against (A) proportional chromosome length (%) and (B) genome size (Mb).

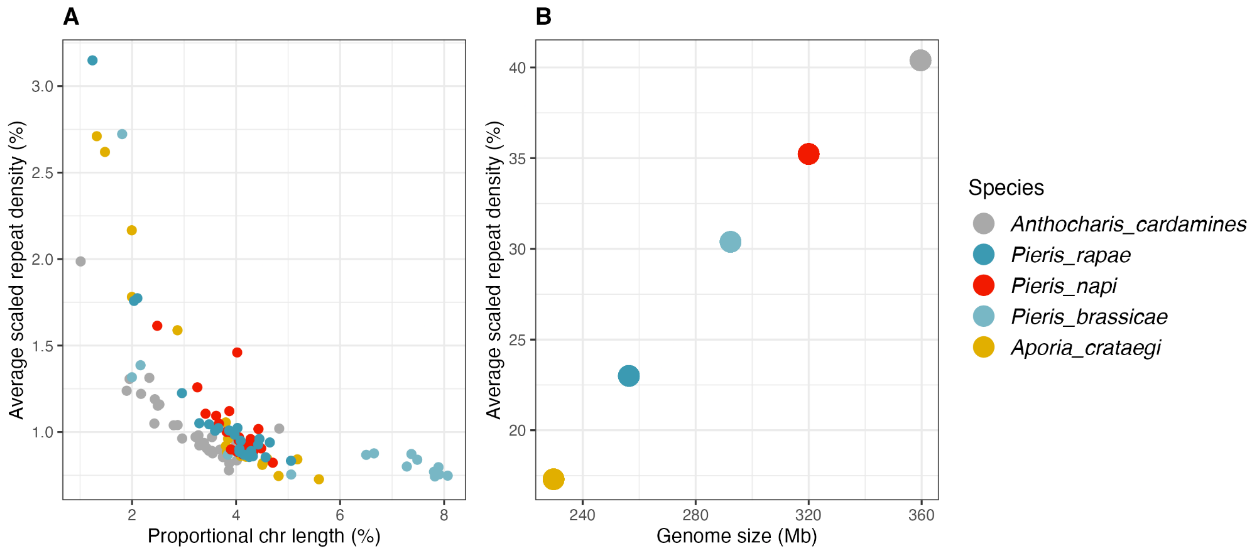


**Fig S30: Relationship between repeat density, genome size and chromosome size in Pierini**

Mean scaled repeat density (%) against (A) proportional chromosome length (%) and (B) genome size (Mb).

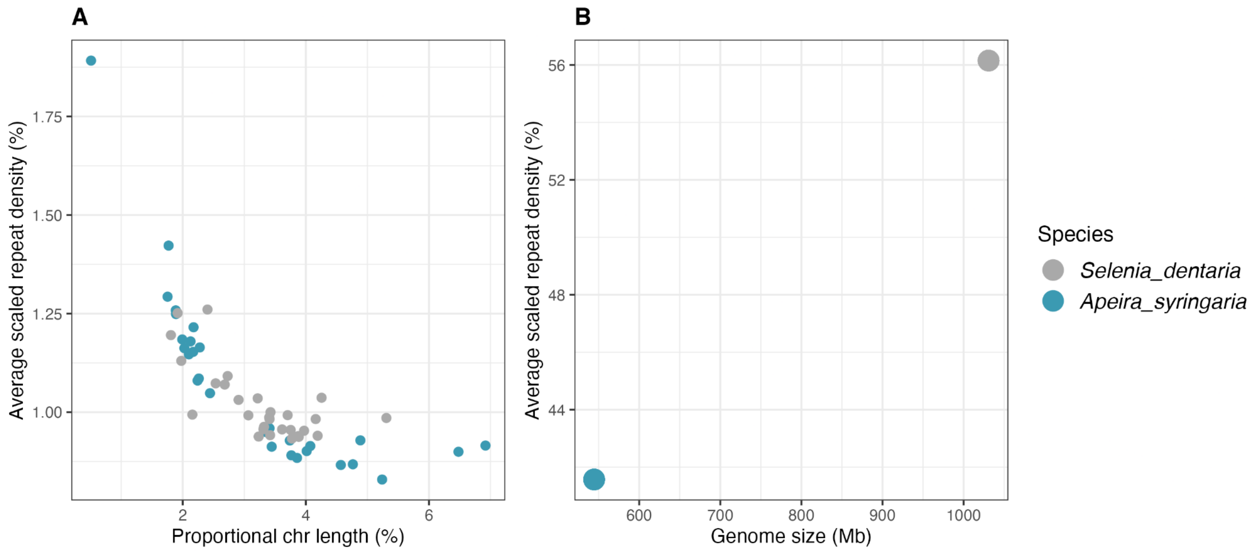


**Fig S31: Relationship between repeat density, genome size and chromosome size in *Apeira***
Mean scaled repeat density (%) against (A) proportional chromosome length (%) and (B) genome size (Mb).

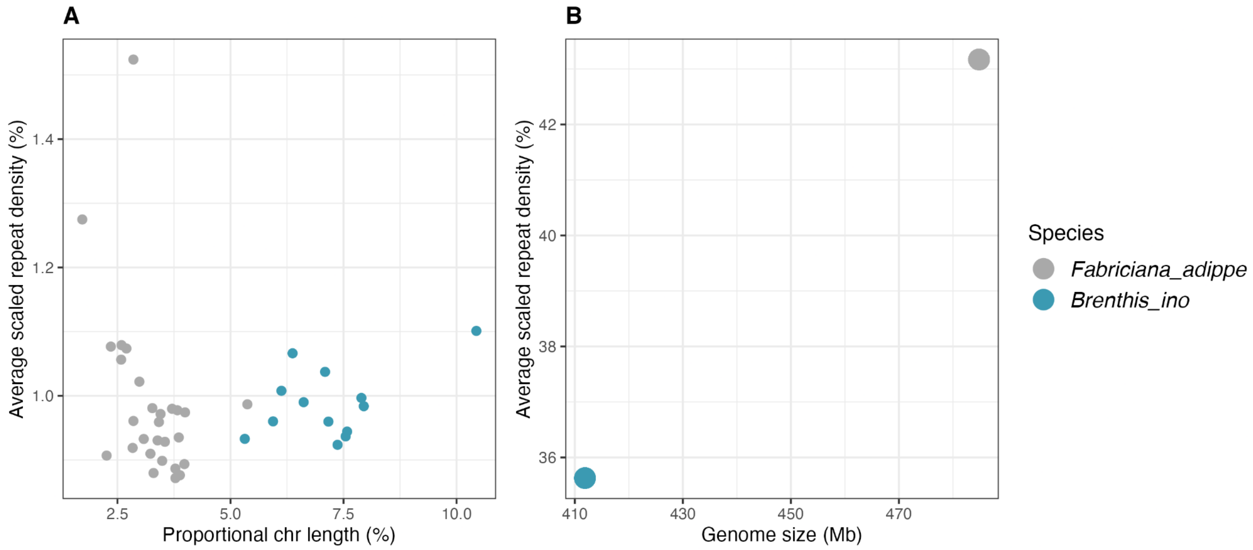


**Fig S32: Relationship between repeat density, genome size and chromosome size in *Brenthis***

Mean scaled repeat density (%) against (A) proportional chromosome length (%) and (B) genome size (Mb).

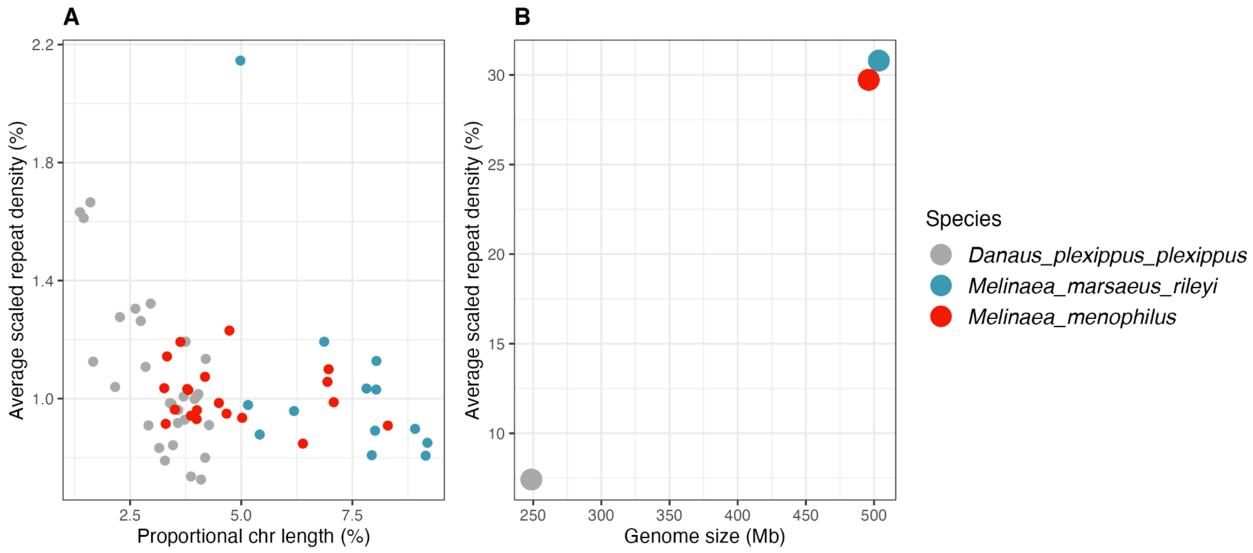


**Fig S33: Relationship between repeat density, genome size and chromosome size in *Melinaea***

Mean scaled repeat density (%) against (A) proportional chromosome length (%) and (B) genome size (Mb).
